# A Versatile, Biocompatible Additive for Spatiotemporal Volumetric Photografting of Biochemical and Mechanical Cues in Diverse Hydrogel Matrices

**DOI:** 10.64898/2026.01.20.700570

**Authors:** Marc Falandt, Camille Bonhomme, Sammy Florczak, Tina Vermonden, Paulina Nunez Bernal, Riccardo Levato

## Abstract

Engineering functional tissue constructs requires not only replicating their 3D architecture but also capturing their dynamic biochemical and mechanical environments. While 3D bioprinting technologies enable spatial control over cell and biomaterial deposition, post-fabrication modulation of material properties remains limited. Photografting approaches allow for spatiotemporal functionalization of certain 3D matrices by chemically binding bioactive factors onto spatially determined regions of a material, but current methods often rely on specialized chemistries with narrow material compatibility. Here, we introduce AddGraft, a biocompatible, off-the-shelf additive designed for semi-orthogonal thiol-ene photografting in vinyl-functionalized hydrogels. AddGraft, a heterobifunctional polyethylene glycol, carries an acrylate moiety for network incorporation during photocrosslinking and a norbornene group for post-crosslinking functionalization. AddGraft integrates into the polymer network during gel crosslinking without altering bulk mechanics, enabling precise modification at any time post-fabrication. We demonstrate compatibility with multiple acrylated biomaterial platforms and light-based volumetric photopatterning technology. Photopatterning achieves high spatial resolution and gradient formation in 3D, while grafting of multi-thiolated crosslinkers allows localized stiffening of hydrogels. Encapsulated human mesenchymal stromal cells exhibit high viability and undergo morphological changes in response to the dynamic tuning of their microenvironment. By decoupling structural and functional roles, AddGraft enables on-demand spatial and temporal control over hydrogel properties. This approach expands the biofabrication toolkit for engineering cell-instructive, 4D tissue environments with translational relevance in regenerative medicine.

## Introduction

A major challenge in tissue engineering lies in recapitulating the intricate spatio-temporal changes in the 3D environment of tissues and organs. During native development, the timely expression and 3D localization of bioactive molecules (*i.e.* growth factors) and dynamic changes in the mechanical environment surrounding cells are crucial in giving rise to tissue-specific functionality.^1–5^ Recently, 3D bioprinting approaches have been explored to create structurally complex cell-laden structures that mimic the cellular organization of the tissue of interest. While a wide library of biomaterials have proven to be compatible with bioprinting techniques, achieving time-dependent, on-demand modifications of their chemical and mechanical properties remain particularly challenging.^1–3^ To overcome the aforementioned challenges in current tissue engineering approaches, the process of photografting bioactive compounds constitutes a promising avenue.^4–9^ Photografting involves the use of light patterns that trigger the chemical immobilization or activation of (bio)molecules inside a polymer network into intricate shapes. To date, this process has been performed through various methods, such as 2D projections via photomasks,^2,4–8^ multiphoton patterning,^2,9–12^ and volumetric photografting.^2,13,14^ When using photomasks, biomolecules can be selectively patterned or activated into two-dimensional shapes, but lack complexity in the z-direction.^4–6^ For enhanced 3D control over the localization of the desired molecules, multiphoton patterning has enabled the creation of micrometer-scale features in 3D planes.^2,4,8,9,12^ However, this technique is limited by the low penetration depth (typically up to 500 µm), far from clinically-relevant scales.^16,17^ In contrast, volumetric photografting allows for large-scale photopatterning within thick polymer structures, creating complex 3D patterns of bioactive molecules within centimeter scale objects in seconds, creating opportunities for the dynamic modification of large tissue mimics.^13,14^ In previous work, thiol-ene click chemistry, based on gelatin norbornene (GelNOR)^13^ and norbornene-functionalized polyvinyl alcohol^14^ have been used to covalently tether thiolated compounds to an already crosslinked hydrogel network. However, this process lacks control over the precise amount of compound that can be photografted, as a variable portion of the norbornene moieties is used to crosslink the hydrogel itself. Novel approaches that decouple hydrogel crosslinking from photografting reactions are therefore necessary. Photografting processes can be achieved through a wide range of chemical reactions.^2,10,13,15–18^ However, this often requires the synthesis of new polymers compatible with the desired photopatterning process, or the direct chemical modification of every protein/compound that is to be grafted.^2,19^ These approaches, while effective, involve complex synthetic steps and require re-optimization depending on the desired material platform and grafted compound(s), limiting the versatility of the process, as well as the chemical scalability of these materials for use in tissue engineering. In this study, we present a novel additive (AddGraft) to enable the (volumetric) photografting process in any (meth)acrylated (bio)polymer. AddGraft consists of a low molecular weight linear polyethylene glycol (PEG) functionalized with an acrylate group on one end and a norbornene on the other (**Figure 1**), and is compatible with various classes of widely used chain growth polymerizable (meth)acrylate-modified biomaterials, including protein-based (*i.e.* Gelatin Methacryloyl, GelMA), polysaccharides (*i.e.* Alginate Methacrylate, AlgMA), or synthetic materials (*i.e.* Polyethylene glycol diacrylate, PEGDA). Since the employed photografting process relies on thiol-ene click chemistry, in absence of accessible thiolated compounds in the pre-hydrogel mixture, no loss of photografting sites is observed. This versatile, off-the-shelf additive, opens new possibilities in the production of dynamic cell culture environments and biofabricated constructs that can undergo time-dependent, spatial modifications throughout their maturation period, potentially mimicking complex developmental changes to bring about advanced function and biomimicry.

**Figure 1.**
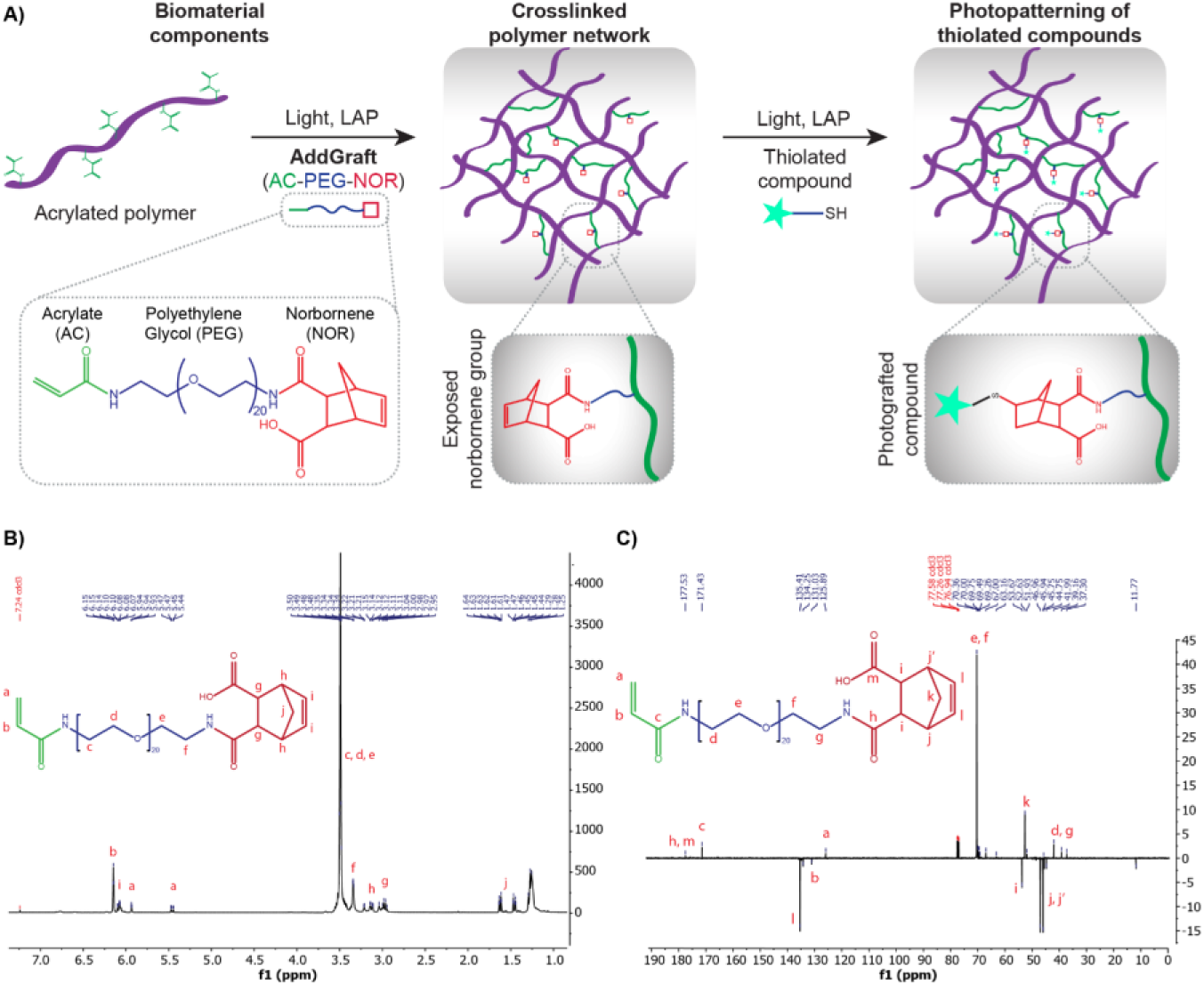
Integration of AddGraft into acrylated polymer networks. A) Schematic illustration of the AddGraft molecule and its incorporation into acrylated biomaterial components and subsequent functionalization of free norbornene moieties with thiolated compounds. C) ^1^H NMR spectrum of AddGraft in deuterated chloroform (CDCl3) with peak assignments showing their corresponding proton on the final structure. D) ^13^C NMR spectrum of AddGraft in CDCl3 with corresponding peak annotations.

## Results and discussion

A mono-protected Boc-PEG_1kDa_-NH_2_ served as the starting material for the synthesis of the AddGraft additive (**Supporting Figure 1**). To introduce the acrylate moiety, the mono-protected PEG was reacted with acryloyl chloride, yielding a mono-acrylated PEG derivative bearing a remaining Boc-protected amine (**Supporting Figure 2**). Subsequently, deprotection of the Boc group using trifluoroacetic acid (TFA) afforded a mono-acrylated PEG with a free primary amine (**Supporting Figure 3**). Lastly, the primary amine was functionalized using carbic anhydride, resulting in the addition of the norbornene moiety on the PEG, yielding the AddGraft final additive. A linear PEG was chosen to be the base of AddGraft given its high-water solubility and widespread availability (**Figure 1A**). ^20–23^ In this proposed setup, the acrylate moiety will enable the crosslinking of AddGraft into the backbone of any acryloyl-based polymer using light-assisted radical polymerization. The crosslinking of the acrylate into a hydrogel network does not interact in any way with the norbornene moieties of the AddGraft, providing a free reactive group after the crosslinking process in known quantities. Once the polymer structure is formed, the exposed norbornene moieties can be used for light mediated thiol-ene click chemistry, effectively binding any thiolated compound to AddGraft. After the synthesis of the additive, proton (^1^H) and carbon-13 (^13^C) nuclear magnetic resonance (NMR) spectra were collected to characterize the purity of the compound (**Figure 1B,C**). As AddGraft was found to form highly viscous aqueous solutions, making it difficult to handle in small amounts, the additive is stored at high concentration in dimethyl sulfoxide (DMSO), to achieve AddGraft supplementation at a concentration of 5 µL/mL in hydrogels well with biocompatibility limits.^24^

**Figure 2.**
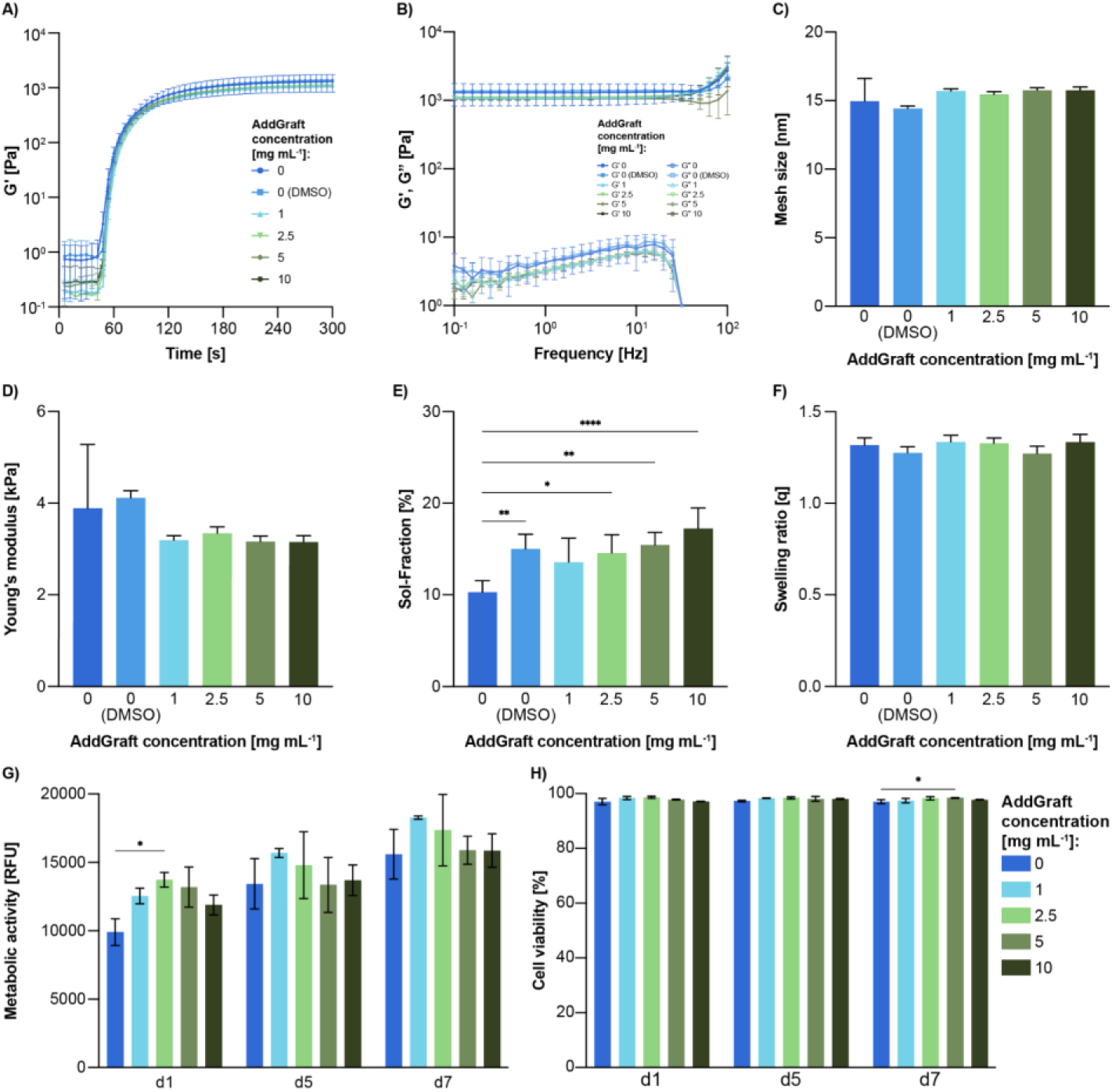
Mechanical, physical and biocompatibility characterization of GelMA hydrogels with varying concentrations of AddGraft. A) Photorheological analysis showing crosslinking kinetics of GelMA formulations with increasing AddGraft concentrations (n = 3). B) Rheological frequency sweeps of GelMA formulations with increasing AddGraft concentrations (n = 3). C) Estimated mesh sizes derived from G′ values in frequency sweep tests (n = 3). D) Compression modulus of hydrogels (n = 5). E) Sol-fraction analysis of hydrogels containing different concentrations of AddGraft (n = 5). F) Swelling ratios of AddGraft-modified hydrogels (n = 3). G) Metabolic activity of hMSCs encapsulated in AddGraft-containing hydrogels over 7 days (n = 3). H) Viability of hMSC-laden hydrogels with varying AddGraft concentrations over a 7-day culture period (n = 3).

**Figure 3.**
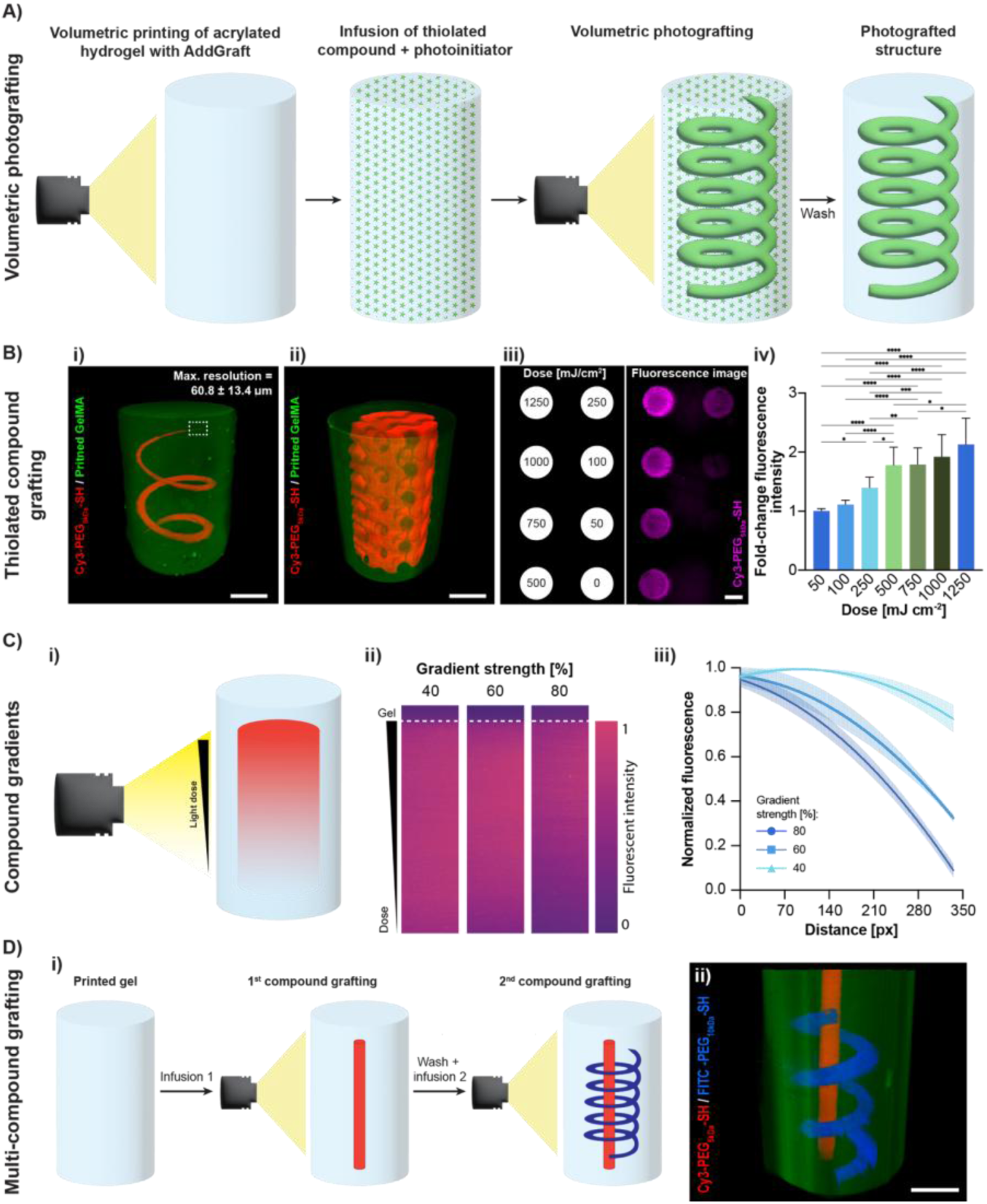
Photografting optimization and characterization of GelMA hydrogels containing 1 mg/mL AddGraft. A) Schematic overview of the photografting process starting with photocrosslinking of the hydrogel construct, infusion with a thiolated fluorescent dye, volumetric photografting, followed by washing to remove ungrafted dye, yielding the photografted hydrogel. B) 3D rendered images from lightsheet microscopy of hydrogels photografted with Cy3-PEG₅ₖDa-SH: i) Thinning spiral pattern used to determine the maximum photografting resolution with AddGraft (scalebar = 2 mm), ii) Complex gyroidal pattern photografted onto the hydrogel (scalebar = 2 mm), iii) Dose-response tests to evaluate photografting efficiency (scalebar = 500 µm), iv) Quantification of fold change in fluorescence intensity at various light doses, comparing grafted areas to background signal (n = 3, N = 4). C) Gradient compound photografting using controlled light intensity gradients: i) Schematic of the gradient photografting setup, ii) Lightsheet images showing varying gradient strengths and corresponding fluorescence intensities from bound molecules, iii) Quantification of normalized fluorescence intensity across the hydrogel (n = 3). D) Multi-compound photografting: i) Schematic illustrating sequential photografting steps with different compounds, ii) 3D lightsheet reconstruction of a multi-compound hydrogel photografted with Cy3-PEG₅ₖDa-SH (red) and FITC-PEG₁₀ₖDa-SH (blue).

Next, we investigated whether the inclusion of AddGraft altered the mechanical and physical characteristics of the formed polymer networks, since its inclusion in the pre-polymer mix entails the availability of additional reactive acrylate groups. Extensive mechanical analysis was performed, using GelMA (gold standard material in bioprinting) crosslinked in the presence of increasing amounts of the additive and DMSO alone. First, the photocrosslinking kinetics of 5% w/v GelMA upon light activated lithium phenyl (2,4,6-trimethylbenzoyl) phosphinate (LAP), was studied. This showed that increasing concentrations of AddGraft (1, 2.5, 5, or 10 mg/mL) did not lead to changes in the crosslinking kinetics and neither did the addition of the base volume of DMSO (0 mg/mL; all experimental conditions contained the same volume of AddGraft/DMSO) (**Figure 2A**). Furthermore, the observed storage modulus plateau of the pristine, additive-free material (1296 ± 378 Pa) did not show significant differences to GelMA supplemented with different AddGraft concentrations (1065 ± 27 Pa, 1115 ± 38 Pa, 1054 ± 34 Pa, 1051 ± 37 Pa, for 1, 2.5, 5, 10 mg/mL AddGraft respectively). Interestingly, gels containing DMSO alone (0 mg/mL) exhibited a significant increase in storage modulus immediately after crosslinking (1372 ± 43 Pa). Although the underlying mechanism is not fully clear, DMSO has been reported to transiently influence polymer network organisation and swelling behaviour of several materials,^25–28^ which may contribute to this effect. Next, a frequency sweep analysis revealed that the storage modulus (G′) and loss modulus (G″) exhibited the same magnitude and frequency dependence in AddGraft-containing gels as in controls (**Figure 2B**). This demonstrates that AddGraft’s incorporation does not alter the gel’s viscoelastic behaviour. Both the photorheological measurements and the frequency sweeps were performed in the viscoelastic region of all the hydrogel formulations as proven with amplitude sweep measurements (**Supporting Figure 4**). The mesh sizes of the crosslinked hydrogels with and without AddGraft were estimated from the G’ values of the frequency sweeps as previously reported.^29^ AddGraft conditions exhibited slightly larger mesh sizes (1 mg/mL: 15.69 ± 0.13 nm, 2.5 mg/mL: 15.46 ± 0.17 nm, 5 mg/mL: 15.75 ± 0.17 nm, 10 mg/mL: 15.76 ± 0.19 nm) than both pristine (14.96 ± 1.34 nm) and 0 mg/mL (14.42 ± 0.15 nm) gels, although not statistically significant. This is consistent with the previously reported lower storage modulus observed post crosslinking. (**Figure 2C**). Likewise, no significant differences were observed in the compressive modulus of hydrogels with different AddGraft concentrations (3.29 ± 0.12 kPa, 3.59 ± 0.22 kPa, 3.59 ± 0.31 kPa, 3.15 ± 0.15 kPa, 3.33 ± 0.37 kPa, for 0, 1, 2.5, 5, and 10 mg/mL AddGraft respectively) compared to the bulk mechanics of the base hydrogel (3.67 ± 0.26 kPa) (**Figure 2D**). The increase in G′ previously observed for the DMSO-only condition during photorheology was no longer present after overnight soaking of the crosslinked samples and subsequent dynamic mechanical analysis, suggesting that DMSO likely diffuses out of the gels during equilibration, eliminating its transient effect on the polymer network. To further evaluate potential differences in network formation, we next quantified the soluble fraction of the gels. Pristine GelMA (10.28 ± 1.12 %) exhibited a significantly lower sol-fraction than GelMA with DMSO (15.02 ± 1.43%) and GelMA with AddGraft concentrations above 1 mg/mL (14.55 ± 1.81%, 15.43 ± 1.24%, 17.24 ± 2.01%, for 2.5, 5, and 10 mg/mL AddGraft respectively), while no significant differences were observed between the DMSO only and AddGraft groups (**Figure 2E**). Furthermore, no significant differences were observed in respect to the swelling ratio of the hydrogels (∼1.3 for all conditions) after crosslinking (**Figure 2F**), nor in the degradation kinetics upon exposure to collagenase type II (**Supporting Figure 5**). Overall, AddGraft does not significantly affect hydrogel mechanics, allowing its concentration, and thus the number of available photografting sites, to be tuned without compromising stability. However, considering the use of DMSO as diluent for the AddGraft stock solution, potential effects of this solvent on biocompatibility were also investigated, by encapsulating human bone marrow derived mesenchymal stromal cells (hMSCs) in GelMA-AddGraft hydrogels. Over a culture period of 7 days, the addition of AddGraft dissolved in DMSO did not affect the metabolic activity of the cells when compared to pristine GelMA samples (**Figure 2G**). Moreover, it was observed that the viability of the hMSCs maintained in all conditions, showing high viability values of > 95% (**Figure 2H**), confirming the safe implementation of this compound for tissue culture applications.

**Figure 4.**
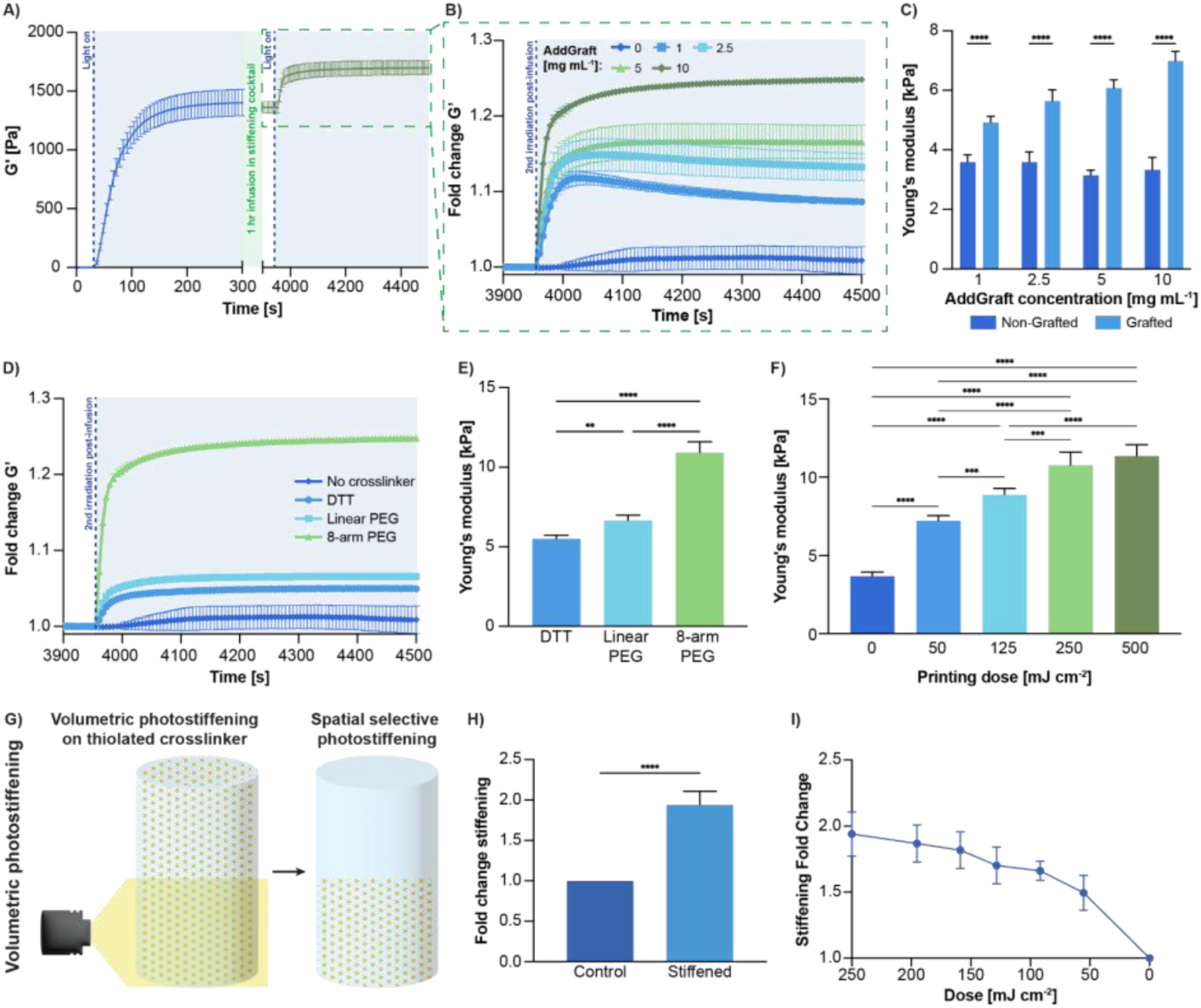
Mechanical characterization of 4D stiffening in AddGraft-infused GelMA hydrogels under varying concentrations, crosslinkers, and light exposure conditions. A) Photorheological assessment showing initial photocrosslinking of GelMA and the subsequent stiffness change following 1 hour infusion with thiolated 8-arm-PEG10kDa-SH crosslinker and a second photoirradiation step (n = 3). B) Fold change in storage modulus of hydrogels with increasing AddGraft concentrations under photorheological testing (n = 3). C) Compression modulus measurements of GelMA hydrogels with increasing concentrations of AddGraft, infused and stiffened using an 8-arm-PEG10kDa-SH (equimolar thiol:norbornene ratio) and LAP photoinitiator (n = 5). D) Photorheological stiffening following 1-hour infusion with various thiolated crosslinkers at 10 mg/mL AddGraft concentration (n = 3). E) Compression modulus analysis of 10 mg/mL AddGraft-infused GelMA hydrogels crosslinked with different thiolated agents: DTT, linear SH-PEG2kDa-SH, and 8-arm-PEG10kDa-SH, all at equimolar thiol:norbornene ratios (n = 3). F) Volumetric photostiffening of 10 mg/mL AddGraft hydrogels under varying light doses, with resulting stiffness assessed by compression modulus (n = 5). G) Schematic illustration of spatially selective volumetric photostiffening using tomographic 3D printing, enabling precise modulation of hydrogel stiffness. H) Fold change in compression modulus measured using a 2 mm indenter on a 10 mg/mL AddGraft hydrogel photostiffened only in the bottom half (n = 3). I) Fold change in stiffness in response to varying light exposure (90% intensity gradient), demonstrating the creation of spatial stiffness gradients within a single 10 mg/mL AddGraft-infused hydrogel construct (n = 3).

**Figure 5.**
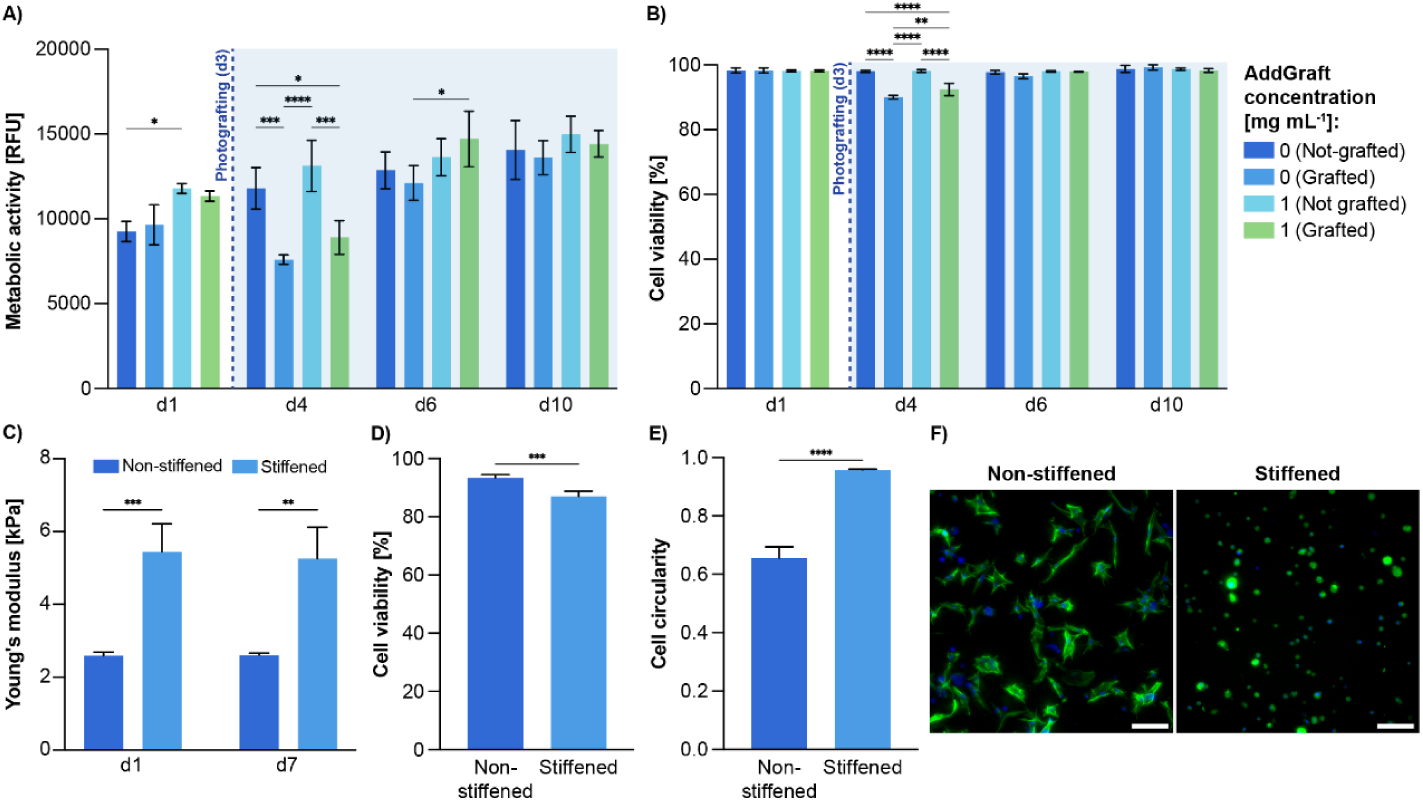
Biocompatibility of the photografting process using AddGraft in GelMA hydrogels with embedded hMSCs. A) Metabolic activity of hMSCs over a 10-day culture period, with photografting of cysteine amino acids performed on day 3 (1 mg/mL AddGraft used to assess procedural biocompatibility) (n = 3). B) Cell viability during the 10-day culture period in AddGraft-containing hydrogels with cysteine photografting at day 3, demonstrating compatibility of the photografting process (n = 3). C) Compression modulus of cell-laden hydrogels on day 1 and day 7, showing maintenance of mechanical integrity after photostiffening and extended culture (photostiffening performed using 10 mg/mL AddGraft in a lower-methacrylation GelMA to maintain cell-tolerant stiffness levels) (n = 3). D) Cell viability on day 7 comparing photostiffened and non-stiffened (control) hydrogels (n = 4). E) Quantification of cell circularity on day 7 in photostiffened versus control hydrogels, indicating morphological differences due to mechanical changes (n = 4). F) Representative fluorescence microscopy images of hMSCs within photostiffened and control hydrogels on day 7 (scalebar = 100 µm).

Having established that AddGraft is biocompatible and does not significantly disrupt the hydrogel’s mechanical properties, we next evaluated its ability to enable post-crosslinking functionalization through volumetric photografting.^13,14^ Because its norbornene groups remain unreacted during gel crosslinking, AddGraft provides reactive handles that enable spatiotemporally controlled chemical modifications even after the primary hydrogel network is formed.^13^ The volumetric photografting procedure employed here, based on a previous study^13^ consisted of the following steps: 1) Fabrication of AddGraft-containing GelMA samples (via volumetric printing or casting), 2) infusion of samples with a grafting cocktail containing LAP photoinitiator and the desired thiolated compound, 3) volumetric photografting, during which filtered back-projections of 405 nm light are delivered to generate a cumulative dose distribution that activates the predefined 3D region, covalently linking the thiolated compound to the free norbornene groups, and 4) washing to remove excess, ungrafted compound (**Figure 3A**). Here, cylindrical samples containing 1 mg/mL AddGraft were fabricated and infused with a grafting cocktail composed of 0.2% w/v LAP (resulting in a final concentration of 0.1% w/v in the gel) and a fluorescent thiolated dye to enable visualization of the grafted structures (0.06% w/v Cy3-PEG₅ₖDa-SH or 0.12% w/v FITC-PEG₁₀ₖDa-SH). Importantly, the system was optimized to balance reactivity with hydrogel integrity. In a previous study using GelNOR, excessive LAP concentrations generated high radical densities that compromised cell viability during the infusion and grafting steps.^13^ To avoid this, the final LAP concentration used here matched that employed during gel crosslinking. Additionally, we assessed the time-dependent uptake of LAP into the hydrogel to determine the minimal infusion time required to reach this concentration to reduce stress on embedded cells (∼30 min – 1 hr; **Supporting Figure 6**). This infusion time was also sufficient to allow homogenous diffusion of the fluorescent PEG dye employed here, as the grafted structures obtained using this protocol spanned the whole depth of the hydrogel samples (**Figure 3Bi,ii**; **Supporting Figure 7**). It is important to note however, that sample infusion time might have to be optimized depending on the compound of interest, as different molecular weights and physicochemical properties may affect diffusion speed throughout the crosslinked hydrogel. The optimized grafting cocktail enabled the photografting of high-resolution features (60.8 ± 13.4 µm), comparable to previously reported systems (**Figure 3Bi**).^13^ This highlights AddGraft’s compatibility with high-precision volumetric photopatterning techniques. Because thiol-ene reactions proceed via step-growth chemistry and can be controlled through timed light exposure, AddGraft enables the formation of defined 3D photografting patterns. This allows the creation of highly complex structures within the hydrogel volume (**Figure 3Bii, Supporting Figure 7**). Constructs photografted with Cy3-PEG₅ₖ_Da_-SH showed measurable and tuneable increases in fluorescence compared to background signal in a light dose-dependent manner. A direct proportionality between light intensity and covalent tethering of fluorescent dye within the hydrogel was observed using controlled static light projections (**Figure 3Biii**). As expected, higher light dose irradiation yielded higher fluorescent intensity regions within the hydrogel as a result of the photografting process (**Figure 3Biv**). AddGraft also enabled the generation of controlled chemical gradients within single hydrogel constructs (**Figure 3Ci**). By programming spatial light-dose gradients into the projection (*i.e.*, an 80% gradient corresponds to a decrease from 125 mJ/cm^2^ dose to 20% across the field), we obtained dose-dependent increases in covalently grafted fluorescent dyes. Lightsheet imaging confirmed well-defined fluorescence gradients for the 40%, 60%, and 80% light-dose profiles (**Figure 3Cii**). Relative to the normalized maximum intensity (1.00), the 80% gradient decreased to 0.09, the 60% gradient to 0.32, and the 40% gradient to 0.77 across the construct (**Figure 3Ciii**). These data demonstrate that AddGraft supports predictable, light-programmed biochemical gradients within hydrogels, enabling precise spatial modulation of chemical cues. When extended to the use of bioactive compounds like growth factors, such a feature could be harnessed to direct cell fate and organization and potentially study developmental signalling processes. This spatial control is not only advantageous for single-compound patterning but also important for recreating the multifactorial signalling environments characteristic of native tissues. To demonstrate this capability, we evaluated whether AddGraft could support multi-compound photografting (**Figure 3D**). Sequential infusion and grafting of distinct thiolated compounds with different molecular weights (Cy3-PEG₅_kDa_-SH and FITC-PEG₁₀_kDa_-SH) was readily achieved, highlighting the high degree of functionalization control provided by this approach, and the possibility to incorporate compounds of different sizes. In contrast, studies employing materials that are solely functionalized with norbornene moieties rely on residual norbornene groups within the network after crosslinking, which limits the availability of photografting sites for sequential incorporation of compounds.^13,14,30,31^

**Figure 6.**
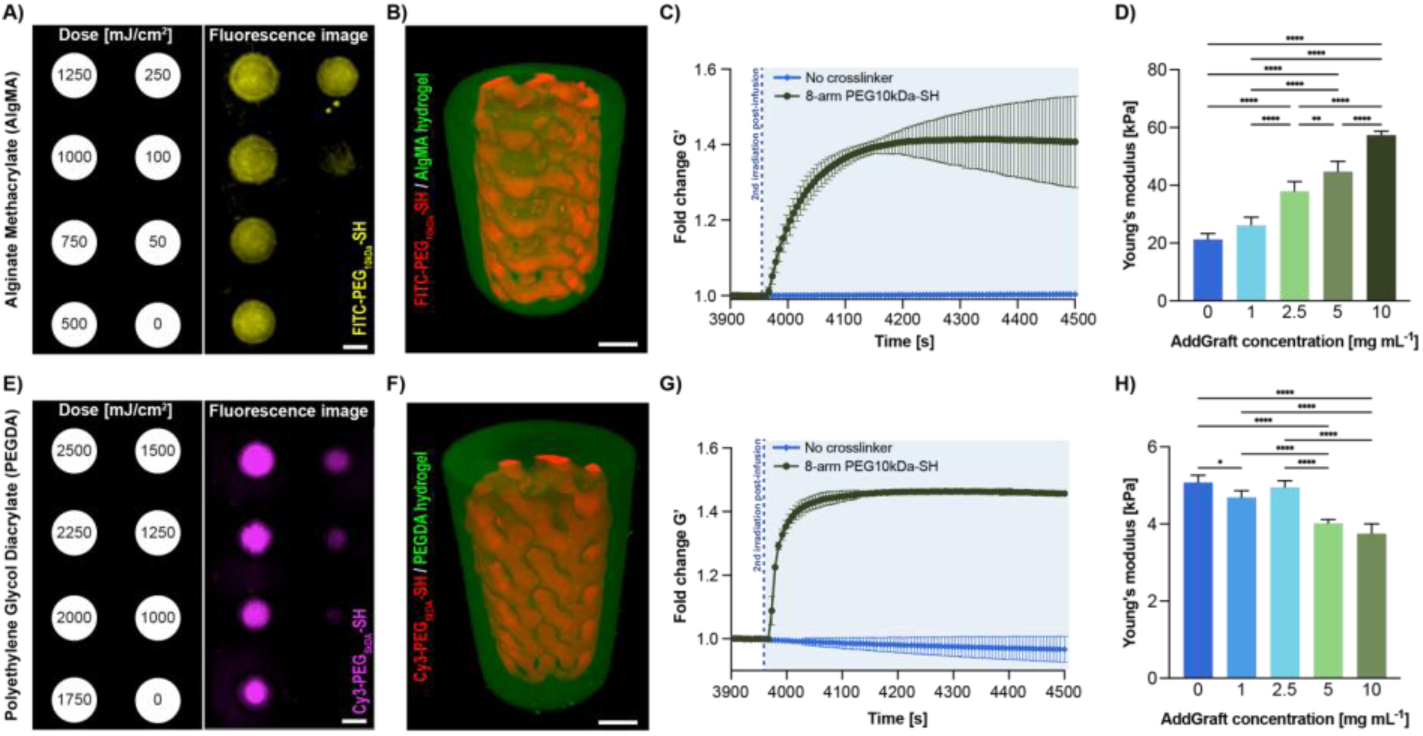
Integration of AddGraft into biomaterials of varying origin, demonstrating photografting optimization, complex patterning, and mechanical modulation in polysaccharide-based and synthetic hydrogel systems. A) Light dose optimization (1000 to 2500 mJ/cm^2^ range) for photografting using FITC-PEG10kDa-SH in AlgMA hydrogels containing AddGraft (n = 2, N = 4) (scalebar = 500 µm). B) Complex photopatterning using fluorescent dye (FITC-PEG10kDa-SH) to demonstrate spatial control of photografting (scalebar = 2 mm). C) Photorheological assessment of stiffening following 1-hour infusion with thiolated crosslinker (n = 3). D) Compression modulus measurements of AlgMA hydrogels with increasing AddGraft concentrations, crosslinked at equimolar thiol:norbornene ratios with 8-arm PEG10kDa-SH (n = 5). E) Light dose optimization (1000 to 2500 mJ/cm^2^ range) for photografting using Cy3-PEG5kDa-SH in PEGDA hydrogels containing AddGraft with 0.008 w/v% TEMPO (n = 3, N = 4) (scalebar = 500 µm). F) Complex photopatterning using fluorescent dye (Cy3-PEG5kDa-SH) to demonstrate spatial control of photografting (scalebar = 2 mm). C) Photorheological assessment of stiffening following 1-hour infusion with thiolated crosslinker (n = 2). D) Compressive modulus measurements of PEGDA hydrogels with increasing AddGraft concentrations, crosslinked at equimolar thiol:norbornene ratios with 8-arm PEG10kDa-SH (n = 5).

Beyond enabling covalent tethering of mono-thiolated dyes and biomolecules, AddGraft can also be used to locally modify the mechanical properties of 3D hydrogel constructs by photografting multi-thiolated crosslinkers. To explore this, different concentrations of AddGraft (1, 2.5, 5, or 10 mg/mL) were incorporated into the hydrogel precursor solution. The formed hydrogels were subsequently stiffened using an 8-arm-PEG_10kDa_-SH crosslinker. To confirm that the initial GelMA crosslinking process was complete prior to the photografting process, photorheological measurements were conducted (**Figure 4A**). For all AddGraft concentrations tested, the storage modulus reached a stable plateau, indicating complete initial network formation before any post-processing modifications. Following crosslinking, the hydrogels were infused with a grafting cocktail containing an equimolar concentration of thiols present in 8-arm-PEG_10kDa_-SH to the corresponding norbornene concentration of each AddGraft condition and 0.2 w/v% LAP for one hour to ensure thorough diffusion into the matrix. Subsequently, a second photorheological measurement was then performed, with a second step of light irradiation. A significant secondary increase in G’ from the initial crosslinking plateau as a result of photostiffening was observed for all AddGraft concentrations (fold change in storage modulus 1 mg/mL: 1.09 ± 0.004, 2.5 mg/mL: 1.13 ± 0.01, 5 mg/mL: 1.17 ± 0.02, 10 mg/mL: 1.25 ±0.002; **Figure 4B**). This concentration-dependent increase in the GelMA storage moduli confirms that AddGraft enables effective post-crosslinking stiffening, while no such stiffening effect was observed in control GelMA hydrogels without AddGraft, even after infusion with the crosslinker and LAP, and the subsequent irradiation step. Additionally, to study the effect of light induced stiffening on the bulk mechanics of the hydrogels, dynamic mechanical analysis (DMA) showed a significant increase in the compression modulus (1 mg/mL: 0.50 kPa, 2.5 mg/mL: 0.57 kPa, 5 mg/mL: 1.47 kPa, 10 mg/mL: 2.81 kPa) of the GelMA hydrogels only when AddGraft was present (**Figure 4C**). While modifying AddGraft concentration is an effective strategy for controlling stiffness, the system also supports the use of various crosslinkers to achieve mechanical tunability independently of total norbornene levels. At the highest tested AddGraft concentration (10 mg/mL), three different crosslinkers were evaluated: dithiothreitol (DTT), linear SH-PEG_2kDa_-SH, and 8-arm-PEG_10kDa_-SH. Employing the same photorheological set-up, we also observed significant crosslinker-dependent fold increases in storage modulus (DTT: 1.05 ± 0.003, SH-PEG_2kDa_-SH: 1.07 ± 0.0002 kPa, 8-arm PEG_10kDa_-SH: 1.25 ± 0.002) aligning with established literature (**Figure 4D**).^13,32^ In short, the addition of 8-arm-PEG_10kDa_-SH in the polymer network results in the incorporation of 4x more junctions than the two linear crosslinkers, which creates a denser, stiffer network post-stiffening. While the incorporation of the linear SH-PEG_2kDa_-SH and DTT should result in similar stiffnesses, as they are both bifunctional compounds, a significantly lower stiffening is observed for DTT. This is likely due to the fact that DTT is the shorter of the compounds and is thus more likely to form intramolecular crosslinks on the same gelatin chain, creating more ineffective crosslinks. Corroborating these photorheological observations, bulk mechanical analysis of hydrogels stiffened with the different crosslinkers revealed a significant increase in compressive modulus compared to the non-stiffened control, as well as distinct differences between crosslinker types (Blank: 3.68 ± 0.23 kPa, DTT: 5.49 ± 0.22 kPa, SH-PEG_2kDa_-SH: 6.65 ± 0.30 kPa, 8-arm-PEG_10kDa_-SH: 10.92 ± 0.61 kPa), further highlighting the versatility of AddGraft-enabled mechanical modulation (**Figure 4E**). To elucidate another approach to modulate the degree of stiffening throughout hydrogel samples, the dose-dependency of the thiolated crosslinkers was also evaluated. Different volumetric light doses (0, 50, 125, 250, and 500 mJ/cm²) were delivered to GelMA samples. Remarkably, even a 50 mJ/cm² light dose (5 seconds) was sufficient to significantly stiffen the hydrogel, while at doses above 250 mJ/cm², a stiffness plateau was reached, highlighting the speed of this process (**Figure 4F**). This maximum stiffness achieved with the volumetric printing set-up (405 nm laser; 11.34 ± 0.73 kPa) was found to be in line with fully UV cured hydrogels. This light dose optimization laid the groundwork for spatially selective mechanical tuning using tomographic printing. In a different experimental setup, only the bottom half of a pre-crosslinked hydrogel construct containing AddGraft was exposed to a 250 mJ/cm² light dose **(Figure 4G**). The result was a localized 2-fold increase in compression modulus of the photostiffened region, confirmed via indentation testing of both irradiated and non-irradiated areas within the same hydrogel (**Figure 4H**). This demonstrates the high spatial resolution and control enabled by AddGraft for inducing heterogeneous mechanical properties within a single bioprinted construct, without the need to perform multi-material printing. Finally, we also demonstrated the possibility of creating mechanical gradients, using a 90% light intensity gradient (highest light dose = 250 mJ/cm^2^) (**Figure 4I**). This resulted in a continuous, linear gradient of stiffness across the hydrogel volume, and opens the door to the generation of mechanically graded environments, where subtle differences in compressive properties could be exploited to modulate cell morphology and behavior. Such capability is a powerful addition to the hydrogel engineering toolkit, allowing for the creation of biomimetic, spatially dynamic constructs that better replicate the mechanical heterogeneity of native tissues, further reinforcing AddGraft’s potential as a key enabler of 4D hydrogel engineering.

With these examples of AddGraft’s potential for (bio)chemical and mechanical patterning established, we next examined the biocompatibility of the photografting process. This evaluation was critical, as volumetric photografting has not yet been demonstrated in cell-laden hydrogels.^13,14^ To specifically assess the effects of the photografting procedure itself (*i.e.* infusion cocktail components and infusion time and a secondary irradiation step), rather than AddGraft concentration, we selected the 1 mg/mL condition as a conservative concentration. Importantly, **Figure 2G,H** already demonstrated the AddGraft dose alone was well-tolerated by cells across the concentration range presented in this study. Mesenchymal stromal cell-laden hydrogels were cultured for three days and then infused for one hour at 37 °C with a cysteine/LAP solution. A second light-irradiation step was applied to graft cysteine onto the AddGraft moieties. Cysteine served as an inert model thiol, enabling evaluation of the infusion and irradiation procedures without introducing functional biological effects. An initial, significant reduction in metabolic activity (irradiated without Addgraft, from 100 to 64.4%; irradiated in the presence of AddGraft: from 100 to 67.8% relative resazurin fluorescence) was observed one day post-photografting (**Figure 5A**). Similarly, cell viability was significantly lower only in samples that received a second step of light irradiation, regardless of the presence of Addgraft (no AddGraft, 90.01 ± 0.32%; AddGraft, 92.43 ± 1.88%), compared to non-irradiated controls that were treated under the same infusion conditions (>98% viability; **Figure 5B**). However, over a subsequent culture period, the irradiated and photografted conditions showed recovery in both viability and metabolic activity, confirming that the photografting process is compatible with viable long-term cell culture. Together, these results indicate that the secondary grafting step – in essence, a second, brief period of radical generation – induces a mild, transient cellular stress response, correlating with previous observations,^33–36^ but cells retain high overall viability (>90%). To examine how cells respond to mechanically patterned environments, we next used a lower-methacrylation GelMA (60% methacrylated lysines), enabling larger stiffness changes upon grafting while maintaining absolute compression moduli within a cell-tolerant range. To maximize the stiffening response in these softer starting hydrogels (2.1-fold increase in compression modulus; **Figure 5C**), 10 mg/mL AddGraft was used in these experiments. MSC-laden GelMA constructs were infused with 8-arm PEG_10kDa_-SH and LAP and subsequently irradiated to induce photostiffening. Mechanical characterization via DMA revealed a significant increase in compression modulus in AddGraft-containing hydrogels, both on day 1 (2.59 to 5.44 kPa) and day 7 (3.15 to 6.10 kPa) post-stiffening (**Figure 5C**), confirming that the photostiffening strategy remains effective in the presence of cells. The stiffened group showed significantly lower viability than the pristine control (93.34 ± 1.15 % vs. 87.00 ± 1.76%) after a week of culture. This is in line with extensive mechanobiological research showing that increased mechanical properties of cells’ 3D microenvironment can lead to changes in cytoskeletal organization and signalling, and in some cases, lead to reduced proliferation or impaired viability (**Figure 5D**).^37,38^ Notably, viability remained above 70%, consistent with mild cellular stress rather than overt cytotoxicity. Furthermore, this mechanical modulation of the cells’ microenvironment resulted in quantifiable morphological changes of embedded cells. In softer 3D matrices, cells can deform and remodel the surrounding network more easily, enabling stretching and even sprouting to occur. In contrast, stiffer matrices (medium-high stiffness ≥5 kPa) restrict matrix remodelling and increase physical confinement, leading to more rounded cell morphologies.^39,40^ Consistent with this, cells in photostiffened regions exhibited significantly higher circularity than those in the non-stiffened matrix (stiffened: 0.96 ± 0.003, non-stiffened: 0.66 ± 0.03; **Figure 5E**). Representative immunofluorescence staining confirmed the observed differences in cell morphology between conditions, with more elongated cells in the non-stiffened group (**Figure 5F**). These findings demonstrate that AddGraft enables precise, spatiotemporal modulation of biochemical and mechanical cues within living 3D cultures. This controllability provides a basis for constructing microenvironments in which both chemical and mechanical signals can be tuned after cell encapsulation.

Having established that AddGraft can be seamlessly incorporated into protein-based materials such as GelMA, we next assessed its versatility as an additive that is not restricted to a specific hydrogel platform. To this end, AlgMA and PEGDA were selected as representative polysaccharide and synthetic polymer systems to evaluate AddGraft’s behavior and the feasibility of photografting within these networks. First, photorheological measurements confirmed that the incorporation of AddGraft into both materials did not affect their crosslinking kinetics, similar to observations in gelatin-based hydrogels (**Supporting Figure 8**). A dose-dependent grafting assay was then conducted on crosslinked AlgMA and PEGDA samples, using 1 mg/mL AddGraft and infused with a fluorescent dye for visualization. Due to the positive charges of Cy3-PEG_5kDa_-SH, the dye could not be effectively washed out of the negatively charged AlgMA hydrogels, and thus a different fluorescent dye was employed (FITC-PEG_10kDa_-SH,). This underlines how different classes of materials may require small procedural modifications to enable the infusion of selected compounds based on their specific diffusional and surface charge properties. Furthermore, it was observed that photografting on AlgMA and PEGDA hydrogels initially led to photobleaching of the fluorescent dye, which we attribute to differences in light absorption and radical reactivity of their polymer backbones. Compared to GelMA, both PEGDA and AlgMA exhibit lower intrinsic light absorbance and reduced radical scavenging capacity, owing to their chemically inert backbones and lack of UV-absorbing amino acid residues.^41,42^ As a result, we hypothesize that a larger fraction of incident photografting light and generated radicals are available to interact with the employed fluorescent dyes, causing photobleaching to occur more rapidly than in GelMA. To mitigate this phenomenon, 2,2,6,6-tetramethylpiperidinyloxyl (TEMPO), a known radical scavenger, was introduced at 0.008 w/v%, a biocompatible concentration previously reported in literature (**Supporting Figure 9**).^13,43,44^ This effectively prevented photobleaching, and enabled dose-dependent grafting onto both AlgMA and PEGDA samples (**Figure 6A, 6E**). The addition of TEMPO, and consequent radical scavenging, resulted in higher required light doses to achieve effective photopatterning (1000 to 2500 mJ/cm^2^), especially in PEGDA hydrogels (**Figure 6E**). Subsequently, complex 3D gyroidal patterns were successfully photografted onto both AlgMA (1250 mJ/cm^2^) and PEGDA (2000 mJ/cm^2^, with 0.008 w/v% TEMPO) samples, demonstrating the broad material translatability of AddGraft-based compound photopatterning (**Figure 6B, 6F**). Finally, we also sought to confirm the potential of AddGraft to mechanically tune these alternative biomaterial sources. Photorheological measurements showed that after a complete photocrosslinking step, both AlgMA (1.41 ± 0.10 fold G’ increase) and PEGDA (1.46 ± 0.01 fold G’ increase) underwent significant mechanical stiffening after infusion in 8-arm PEG_10kDa_-SH and a second round of irradiation **Figure 6C, 6G**). For AlgMA hydrogels, photostiffening with different AddGraft concentrations resulted in significant increases in compression modulus with increasing AddGraft concentrations (26.14 ± 2.53 kPa, 37.95 ± 3.01 kPa, 44.76 ± 3.17 kPa, 57.35 ± 1.28 kPa, for 1, 2.5, 5, and 10 mg/mL AddGraft respectively) compared to the base material (21.26 ± 1.82 kPa). (**Figure 6D**). Moreover, conventional stiffening of AlgMA using calcium chloride (CaCl_2_) was still possible after AddGraft-based photostiffening, expanding the mechanical tuneability of this material platform, and reaching stiffnesses of approximately 100 kPa (**Supporting Figure 10**).^45–47^ Furthermore, higher AddGraft concentrations led to reduced hydrogel shrinkage (**Supporting Figure 11**), indicating that comparable stiffness could be achieved with less dimensional change compared to AddGraft-free controls. As for AddGraft-containing PEGDA hydrogels, while immediate photostiffening was shown through photorheological measurements (**Figure 6G**), stiffened samples that were equilibrated overnight for dynamic mechanical analysis exhibited lower Young’s moduli than pristine samples. Specifically, a significant decrease in compression modulus was measured with increasing AddGraft concentrations (4.69 ± 0.16 kPa, 4.95 ± 0.15 kPa, 4.02 ± 0.09 kPa, 3.75 ± 0.23 kPa, for 1, 2.5, 5, and 10 mg/mL AddGraft respectively), compared to the base material (5.08 ± 0.16 kPa) (**Figure 6H**). This softening effect was associated with an increased swelling of the polymer upon reaching swelling equilibrium. This effect is likely due to an increase in hydrophilicity of the gels, as AddGraft-stiffening entailed the incorporation of additional PEG molecules onto a hydrophilic backbone, therefore leading to lower mechanical stiffness (**Supporting Figure 12**). ^48,49^ While unexpected, this strategy could have potential for applications in which locally reducing matrix stiffness is desired. Nonetheless, these changes in PEGDA mechanics, and the initial increase in G’ observed in non-swollen hydrogels still demonstrate the successful incorporation of AddGraft-bound crosslinkers. The use of PEG-free crosslinkers could be investigated to induce long-term stiffening of this particular polymer in future studies. Taken together, the ability to incorporate AddGraft into protein-, polysaccharide-, and synthetic polymer-based hydrogels demonstrates that its photografting functionality is not dependent on a specific matrix chemistry. Although different material platforms may require some optimization of the process to accommodate differences in their chemical and physical properties, the core grafting process can be translated reliably across multiple material platforms that are widely used in the field of biofabrication and tissue engineering. This adaptability establishes AddGraft as a broadly applicable additive for spatially controlled hydrogel modification.

## Conclusion

This work introduces a versatile and modular strategy for introducing dynamic, spatially controlled functionality into a broad range of acrylated biomaterials. Using light to drive selective thiol-ene coupling, AddGraft enables user-defined biomolecules or crosslinkers to be covalently attached at chosen locations and time points. In doing so, it decouples the design of the base hydrogel from the incorporation of instructive cues, eliminating the need to modify the underlying biomaterial. We demonstrate that AddGraft supports photografting of fluorescent markers and localized modulation of stiffness with high spatial freedom, facilitated by its compatibility with tomographic volumetric bioprinting. Importantly, this additive functions robustly across chemically distinct hydrogel platforms, including acrylated protein-based, polysaccharide, and synthetic polymer systems, with only minimal system-specific adjustments. When grafting bioactive proteins, however, it is important to consider that thiol-ene chemistry may engage cysteine residues within native domains, potentially reducing activity. Our previous work showed that thiol-ene grafting of VEGF onto gelatin-based hydrogels preserved bioactivity,^13^ but post-grafting functional assays remain essential. Given the reliance on light-mediated reaction, we envision this additive to be relevant for other light-based bioprinting technologies, including multiphoton lithography. Modification of the hydrogel environment was demonstrated to be possible in presence of living cells and well-tolerated. Moreover, the modular nature of AddGraft allows it to be easily adapted for different biochemical factors, or mechanical properties, offering a powerful toolkit for exploring how microenvironmental dynamics shape cell behavior, differentiation, and tissue organization. Given its simplicity, the additive could be readily adopted by multiple laboratories, making spatial modifications of hydrogels broadly accessible. By enabling post-fabrication editing of both biochemical and mechanical cues, AddGraft offers a flexible and scalable platform for designing more physiologically relevant and dynamically tunable 3D culture systems.

## Acknowledgements

This project received funding from the European Research Council (ERC) under the European Union’s Horizon 2020 research and innovation programme (grant agreement No. 949806, VOLUME-BIO). R.L. acknowledges the funding from the Gravitation Program “Materials Driven Regeneration” (024.003.013) and from the Summit program “DRIVE-RM”, funded by the Netherlands Organization for Scientific Research. R.L. also acknowledges financial support from the Dutch Research Council (Vidi, 20387).

## Material and Methods

### Materials

Gelatin from porcine skin (Type A, 175g Bloom, Sigma-Aldrich) was used for hydrogel synthesis. PEGDA (Mw = 8kDa) and sodium alginate from brown algae (12-40 kDa), cellulose dialysis membrane tubes (molecular weight cutoff = 12 kDa) were purchased from Sigma-Aldrich. Lithium phenyl-2,4,6-trimethylbenzoylphosphinate (LAP) was purchased from Tokyo Chemical Industry (Tokyo, Japan). SH-PEG_2kDa_-SH (Mw = 2 kDa), 8-arm PEG_10kDa_-SH (Mw = 10 kDa), Cy3-PEG_5kDa_-SH (Mw = 5 kDa), and FITC-PEG_10kDa_-SH (Mw = 10 kDa) were purchased from Biopharma PEG (Watertown, USA). Boc-PEG_1kDa_-NH_2_ (Mw = 1 kDa) was purchased from Creative PEGWorks (Chapel Hill, USA). All other chemicals or reagents were purchased from Sigma-Aldrich unless stated otherwise.

### Synthesis of AG1

Boc-PEG_1kDa_-NH₂ (1.00 g, 1.0 mmol, 1.0 eq) was dissolved in dry dichloromethane (DCM, 10 mL) at 4°C. Aqueous NaOH (20% w/v, 200 µL, 1.0 mmol, 1.0 eq) was added to the solution. Acryloyl chloride (97 µL, 1.2 mmol, 1.2 eq) was then added dropwise under stirring. The reaction mixture was stirred on ice for 3 hours. Upon completion, the organic layer was separated, washed with brine (3 × 5 mL), dried over anhydrous magnesium sulfate (MgSO₄), filtered, and concentrated under reduced pressure. The resulting crude product was lyophilized to remove residual water, yielding **AG1** as a white solid (∼75% yield). ^1^H NMR (400 MHz, CDCl_3_) δ: 6.26 (d, J = 1.8 Hz, 1H), 6.19 – 6.13 (m, 1H), 5.60 (dd, J = 10.1, 1.8 Hz, 1H), 3.82 – 3.79 (m, 1H), 3.63 (d, J = 3.2 Hz, 94H), 3.54 – 3.49 (m, 4H), 3.47 – 3.44 (m, 0H), 3.29 (d, J = 5.4 Hz, 2H), 1.43 (s, 9H). ^13^C NMR (100 MHz, CDCl_3_) δ: 130.60, 126.08, 70.54, 70.50, 70.23, 70.17, 69.79, 58.61, 39.29, 28.41.

### Synthesis of AG2

**AG1** (1.0 eq) was dissolved in dry DCM (100 mM) at room temperature under nitrogen. The solution was continuously degassed with nitrogen bubbling prevent DCM evaporation during the reaction. Next, trifluoroacetic acid (TFA, equal volume to DCM) was added, and the mixture was stirred for 30 minutes under a constant nitrogen atmosphere. The reaction mixture was then evaporated under a stream of nitrogen and lyophilized to afford **AG2** as a yellow oil (quantitative yield). ^1^H NMR (400 MHz, CDCl_3_) δ: 6.27 – 6.18 (m, 1H), 5.68 (dd, J = 9.8, 1.8 Hz, 2H), 3.82 – 3.73 (m, 2H), 3.70 – 3.42 (m, 80H), 3.13 (q, J = 5.5 Hz, 2H). ^13^C NMR (100 MHz, CDCl_3_) δ: 159.34, 158.94, 158.54, 130.10, 127.53, 116.65, 113.79, 70.38, 70.35, 70.29, 70.26, 70.21, 70.18, 70.05, 70.03, 69.90, 69.87, 69.80, 69.73, 69.62, 69.57, 69.46, 69.26, 67.94, 66.68, 39.82, 39.75, 25.48.

### Synthesis of AddGraft

**AG2** (1.0 eq) was dissolved in dry tetrahydrofuran (THF, 50 mM). Carbic anhydride (1.2 eq) and N,N-diisopropylethylamine (DIPEA, 3.0 eq) were added. The reaction mixture was heated to 60 °C and stirred overnight. Progress was monitored via TLC and ninhydrin. If free amines were still present, an additional equivalent of carbic anhydride and DIPEA was added, and the mixture stirred overnight again. After completion, the solvent was removed under reduced pressure. Water was added to the resulting oil to precipitate excess carbic anhydride. The mixture was filtered, and the product was lyophilized to yield AddGraft as a yellow oil (∼80% yield). ^1^H NMR (400 MHz, CDCl_3_) δ: 6.15-6.14 (s, 1H), 6.10 – 6.07 (m, 2H), 5.94 – 5.93 (d, J = 5.92, 1H), 5.47 – 5.44 (t, J = 2.9, 1H), 3.50 – 3.48 (m, 82H), 3.35 – 3.34 (m, 2H), 3.22 – 3.11 (m, 2H), 3.04 – 2.95 (m, 2H), 1.64 – 1.61 (m 1H), 1.47 – 1.44 (m, 1H). ^13^C NMR (100 MHz, CDCl_3_) δ: 177.53, 171.43, 135.41, 134.25, 131.01, 125.89, 70.36, 70.00, 69.75, 69.49, 69.26, 67.00, 63.16, 53.67, 52.63, 51.93, 46.96, 45.94, 45.75, 44.75, 41.99, 39.16, 37.30.

### Gelatin methacryloyl (GelMA) synthesis

GelMA was synthesized according to previously reported literature.^50^ Briefly, Type A Gelatin was dissolved at 50°C in CB-Buffer (carbonate-bicarbonate buffer pH 9, 0.1 M concentration) at 10 w/v% concentration. To reach the desired degree of methacrylation (DoM), 71 µL of methacrylic anhydride (MAA) per gram of gelatin was used in the reaction. The MAA was added at 6 instances, every 11 minutes. After every addition of MAA the pH of the reaction was stabilized to 9. Exactly 90 minutes after the first addition of MAA, the reaction was stopped by lowering the pH to 7.4. Subsequently, the solution was centrifuged at 4000 rpm at room temperature for 5 minutes. The supernatant was collected and diluted with MilliQ water to reach a concentration of 5 w/v%. The solution is then dialyzed against MilliQ for 5 days at 4°C, sterile filtered and lyophilized to yield the dry product (DoM : 60% and 80%).

### AlgMA synthesis

AlgMA was synthesized based on previous literature.^51^ Briefly, Alginate was dissolved in MilliQ water at a 2 w/v% concentration. Subsequently equivolume of MAA was added to the dissolved alginate solution at room temperature. The pH of the mixture was checked periodically and adjusted to pH 7 using NaOH (5 M). The reaction was allowed to stir for 3 days at room temperature. After 3 days, the solution was poured into a 2.5 fold volume of cold EtOH to precipitate the AlgMA. The precipitate was dried at 40 degrees overnight to yield the dry product. For further purification, the AlgMA was dissolved in MilliQ at a 1 w/v% concentration and dialyzed against MilliQ for 5 days at 4°C. Followed by sterile filtration and lyophilization of the purified AlgMA. Degree of Functionalization (DoF) is calculated according to proton NMR (**Supporting Figure 14**) following **Equation 1**, and **Equation 2**. Calculated DoF for AlgMA ∼30%.

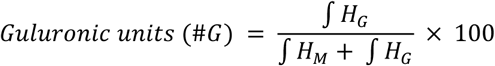

Equation 1. Formula to calculate the amount of guluronic units in the AlgMA. HG = anomeric carbon of guluronic unit (ca. 4.80-5.00 ppm), HM = anomeric carbon of mannuronic unit (ca. 4.50 ppm).

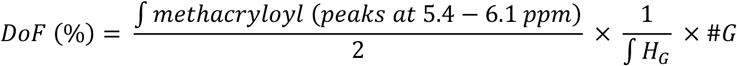

Equation 2. Formula to calculate the Degree of Functionalization (DoF) of methacrylates onto the AlgMA.

### Hydrogel preparation

Unless stated otherwise, all experiments are conducted using GelMA (5 w/v%) supplemented with LAP (0.1 w/v%) in phosphate buffer saline (PBS). For different concentrations of AddGraft, a stock dilution was made so the final volume of DMSO in the hydrogel precursor would never surpass 5 µL/mL of bioink. Concentrations of AddGraft were 0, 1, 2.5, 5, and 10 mg/mL (0, 1, 2.5, 5, and 10 mM of AddGraft). AlgMA hydrogels were prepared at a 3 w/v% concentration. PEGDA (Mw = 8 kDa) hydrogel solution was prepared at a 4 w/v% concentration.

### Photorheology, frequency sweep, amplitude sweep

Photorheology experiments on hydrogel precursor solution with and without AddGraft were assessed using a DHR20 rheometer (TA Instruments, The Netherlands). 100 µL of hydrogel was used with a gap size of 300 µm, using a 20.0 mm EHP stainless steel plate as geometry. Time sweep experiments were performed at 1.0 Hz frequency, with a 1.0% constant strain at 20°C (n = 3). After 30 seconds, the light source was activated (Omnicure S2000 Elite, λ = 405 nm, I = 7.5 – 30 mW/cm^2^) and kept on for the remaining 2.5 minutes of the experiment. Subsequently, a frequency sweep from 0.1 Hz to 100 Hz was performed at a constant 1.0% strain at 20°C. Next, an amplitude sweep was performed on the same sample from 0.1 to 100% strain at a constant 1.0 Hz frequency at 20°C. For the assessment of stiffening with the photorheometer, a second time sweep was performed after the first time sweep experiment without measuring a frequency, or amplitude sweep. For this a specially made immersion bath was filled with a 500 µL thiolated crosslinker infusion solution for 1 hour. Subsequently, the second time sweep experiment was performed at 1.0 Hz frequency, with a 1.0% constant strain at 20°C (n = 3). After 60 seconds, the light source was activated (Omnicure S2000 Elite, λ = 405 nm, I = 30 mW/cm^2^) and kept on for the remainder of the experiment. The mesh size of the crosslinked hydrogels was calculated from the maximum G’ obtained in the frequency sweep as previously reported (Equation 3).^29^

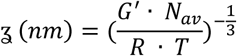

Equation 3. Mesh size formula.

*Mechanical characterization (compression modulus):* Hydrogel precursor solutions were poured into cylindrical molds (Ø 6 mm, 2 mm height) and allowed to crosslink for 10 minutes to ensure full network formation. Following gelation, the constructs were incubated in PBS at 37 °C overnight to allow equilibration to their swollen state. Compressive mechanical testing was then conducted using a dynamic mechanical analyzer (DMA Q850, TA Instruments, The Netherlands). Samples were compressed at a constant strain rate of 20% per minute until reaching 30% strain with a 0.001 N preload force. The compressive modulus was determined from the linear region of the stress–strain curve, specifically within the 10–15% strain range.

*Sol fraction and swelling ratio:* Sol-fraction of AddGraft containing hydrogel formulations were tested according to recent literature.^13^ Briefly, hydrogels with different AddGraft concentrations (0, 1, 2.5, 5, 10 mg/mL) were crosslinked into cylindrical samples (8 mm, 3 mm height). The hydrogels were weighed after crosslinking for their initial mass (Mass_wet,t0_). Next, the hydrogels were lyophilized to yield the dry mass at t = 0 (massdry,t0). The samples were stored in PBS again to ensure swelling of the dry gels and placed in the incubator at 37°C overnight. The wet mass of the hydrogels was measured as masswet,t1. The samples were lyophilized, and the mass of the dry samples was measured as massdry,t1. The soluble fraction (sol-fraction; **Equation. 4**) and swelling ratio (**Equation. 5**) of the hydrogels were then calculated.

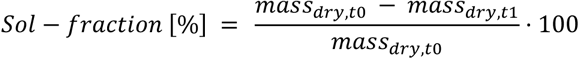

**Equation 4**. Sol-fraction formula for analysis of the crosslinking properties of the hydrogel formulations.

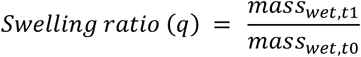

**Equation 5**. Swelling ratio for analysis of the swelling behavior of the hydrogel formulations.

### Metabolic activity and viability of cell-laden AddGraft hydrogels

Cylindrical hMSC-laden constructs (1 × 10^6^ cells mL^−1^; 6 mm diameter × 2 mm height) were casted using GelMA with AddGraft (0, 1 mg/mL) and crosslinking for 8 min in a CL-1000 Ultraviolet Crosslinker (λ = 365 nm; UVP, USA). Samples were cultured in Dulbecco’s Modified Eagle Medium (DMEM, Gibco, 31966, The Netherlands), supplemented with 10 v/v% heat-inactivated fetal bovine serum (FBS, Gibco, The Netherlands), 1 v/v% penicillin-streptomycin (Life Technologies, The Netherlands), l-ascorbic acid-2-phosphate (0.2 × 10–3 M, Sigma-Aldrich, The Netherlands) and 1 µL/mL basic Fibroblast Growth Factor (bFGF, R&D Systems, USA) for 10 days, which was refreshed every two days. Metabolic activity (n = 3) was measured with a resazurin assay (resazurin sodium salt, Alfa Aesar, Germany) after 1, 4, 5, 6, 7 and 10 days. Cell viability was evaluated using a live/dead assay (Calcein, ethidium homodimer, Thermo Fischer Scientific, The Netherlands) after 1, ,4, 5, 6, 7 and 10 days (n = 3), imaged by a Thunder imaging system (Leica Microsystems, Germany).

### Enzymatic degradation

GelMA hydrogels (Ø 6 mm, 2 mm height) were first allowed to swell overnight in PBS. Following this equilibration step, the samples were incubated at 37°C in an enzymatic solution consisting of 0.2 w/v% collagenase type II prepared in Dulbecco’s Modified Eagle Medium (DMEM, Gibco, 31966, The Netherlands), supplemented with 10 v/v% heat-inactivated fetal bovine serum (FBS, Gibco, The Netherlands),1 v/v% penicillin-streptomycin (Life Technologies, The Netherlands) and l-ascorbic acid-2-phosphate (0.2 × 10–3 M, Sigma-Aldrich, The Netherlands). Hydrogel samples were collected from the enzyme solution at predefined time intervals (15, 30, 45, and 60 minutes; n = 3 per time point). To assess the extent of degradation, the wet mass of each sample was recorded and compared to its initial pre-incubation mass.

### Volumetric printing

Prepolymer solutions were transferred into cylindrical borosilicate glass vials (Ø 10 mm) and subsequently placed into a commercial volumetric 3D printer (Tomolite V2, Readily3D, Switzerland), equipped with a 405 nm laser operating at an average light intensity of 8.4 mW/cm² within the build volume. To initiate physical gelation of the gelatin-based formulations, the samples were cooled to 4°C prior to the printing step. Custom STL models were imported into the printer’s control software (Apparite, Readily3D, Switzerland) to define the target geometries. Upon completion of the volumetric printing, the vials were warmed to 37°C, and the constructs were gently rinsed with pre-warmed PBS (37°C) to recover the printed structures.

### Photografting process and infusion

Crosslinked constructs (GelMA, AlgMA, PEGDA), containing AddGraft (1 mg/mL), were subjected to a second printing step to induce spatio-selective photopatterning. The crosslinked hydrogels were infused in a mixture of fluorescent probe molecule, Cy3-PEG_5kDa_-SH (0.06 w/v%) or FITC-PEG_10kDa_-SH (0.12 w/v%), LAP (0.2 w/v%), and with TEMPO (0, 0.004, or 0.008 w/v% concentration) in PBS at a 1 to 1 volume ratio. The hydrogel was allowed to fully infuse with the compounds prior to volumetric photografting (overnight at 4°C for cell free experiments, 1 hour at 37°C degrees for cell-laden hydrogels). The crosslinked and infused hydrogels were placed into a volumetric printing vial and exposed to the volumetric printing process. The constructs were washed with PBS for a maximum of 5 days, until the untethered dye was completely removed from the hydrogel. Subsequently, the photografted structures were imaged with a light-sheet microscope. This process can be repeated to ensure multiple compound grafting in the same hydrogel. For the quantification of grafting intensity at multiple light doses, a built in option of the volumetric printer, named Dose Test, was used. Infused and crosslinked rectangular hydrogels (24 mm length, 9 mm width, 2 mm height) were loaded into quartz cuvettes (CV10Q1400FS, Thorlabs) placed in a specifically designed cuvette holder and transferred into the volumetric printer. The Dose Test function was used to project a variable dose range matrix of dots (Ø 0.8 mm dimensions) onto the hydrogel. The illuminated hydrogels were washed with PBS to remove uncrosslinked fluorescent dye and imaged with a Leica thunder microscope. The grafting intensity of crosslinked fluorescent probes was measured as an increase of fluorescence of the intended region vs the background signal (**Equation 6**).

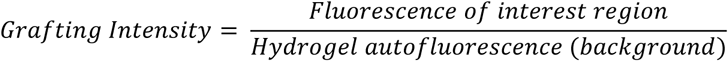

**Equation 6**. Grafting intensity formula for analysis of grafting on AddGraft containing hydrogels.

### Photostiffening infusion and process

Crosslinked AddGraft containing hydrogels (GelMA, AlgMA, PEGDA) were infused with thiolated crosslinkers to enable material photostiffening. Hydrogels were infused with thiolated crosslinkers DTT, SH-PEG_2kDa_-SH, or 8-arm-PEG_10kDa_-SH together with 0.2 w/v% LAP. Thiolated crosslinker concentrations were calculated to provide an equimolar ratio of thiols to norbornene groups based on the nominal norbornene concentration introduced via AddGraft, assuming one norbornene moiety per AddGraft molecule. AddGraft concentration and molecular weight was used to estimate [NOR] (**Equation 7**). The infusion of the hydrogels was done at 4°C overnight to ensure complete diffusion of the thiolated crosslinkers. Subsequently, the infused hydrogels were exposed to UV light (10 minutes) or the volumetric printer (varying light doses) to ensure photostiffening of the construct. Only for AlgMA hydrogels, a second round of stiffening was induced by submerging the already photostiffened hydrogels in a solution of CaCl_2_ (50 mM) for 2 hours. Stiffness of the samples was measured by DMA. For indentation measurements of spatially controlled photostiffened samples, a rectangular hydrogel (7 mm width, 7 mm height, 20mm length) was crosslinked. A 2 mm indenter was used to compress the hydrogel material in selected areas of the construct. Samples were exposed to a constant strain rate of 20% per minute until 30% strain was reached, with a 0.001 N preload force. The modulus was calculated between 10-15% strain of the stress-strain curve.

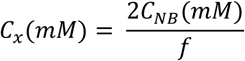

**Equation 7**. Equation to calculate the required thiolated crosslinker concentration to achieve an equimolar norbornene : thiol ratio. CNB = norbornene concentration; *f* = thiols present in crosslinker molecule.

### Biocompatibility assessment of photografting

Cylindrical hMSC-laden constructs (1 × 10^6^ cells mL^−1^; 6 mm diameter × 2 mm height) were casted using GelMA with AddGraft (0, 1 mg/mL) and crosslinking for 8 min in a CL-1000 Ultraviolet Crosslinker (λ = 365 nm; UVP, USA). On day 3 of culture, the cell-laden hydrogels were incubated for 1 hour at 37°C in a mixture containing cysteine (at an equimolar ratio of thiol to AddGraft) and 0.2 w/v% LAP. Subsequently, the samples, excluding the controls, were exposed to a second round of light irradiation for 8 minutes. Metabolic activity and cell viability were assessed as previously described.

### Cell experiment stiffening

Cylindrical hMSC-laden constructs (500 × 10^5^ cells mL^−1^; 6 mm diameter × 2 mm height) were casted using GelMA with AddGraft (0, 10 mg/mL) and crosslinking for 8 min in a CL-1000 Ultraviolet Crosslinker (λ = 365 nm; UVP, USA). Samples were cultured in Dulbecco’s Modified Eagle Medium (DMEM, Gibco, 31966, The Netherlands), supplemented with 10 v/v% heat-inactivated fetal bovine serum (FBS, Gibco, The Netherlands),1 v/v% penicillin-streptomycin (Life Technologies, The Netherlands), l-ascorbic acid-2-phosphate (0.2 × 10–3 M, Sigma-Aldrich, The Netherlands), and 1 ng/mL basic Fibroblast Growth Factor (bFGF, R&D Systems, USA) for 7 days, which was refreshed every two days. On day 0 of culture, the cell-laden hydrogels were incubated for 1 hour at 37°C in a mixture containing 8-arm PEG_10kDa_-SH (at an equimolar ratio of thiol to AddGraft) and 0.2 w/v% LAP. Subsequently, the samples, excluding the controls, were exposed to a second round of light irradiation for 8 minutes. Cell viability (n = 4) and compression modulus (n = 3) were measured on day 1 and day 7 of culture. Cell circularity was measured on day 7 (n = 3).

### LAP diffusion kinetics

For the assessment of the diffusion kinetics of LAP, GelMA hydrogels were crosslinked using UV oven for 10 minutes (Ø 6 mm, 2 mm height) and washed overnight at 37°C in PBS. A calibration curve of LAP in PBS was made at 0, 0.01, 0.025, 0.05, 0.075, 0.10, 0.15, 0.2 w/v% in PBS. Then the GelMA hydrogels were infused with a 1 to 1 volume ratio of a 0.2 w/v% LAP solution and allowed to incubate in the LAP mixture. The hydrogels were taken out of the infusion mix at different timepoints (15, 30, 60, 120 min). The absorbance of the infusion mix of samples at different time points was measured by a CLARIOstar Plus® (BMG Labtech, Germany) plate reader at λ = 405 nm (n = 3).

### Statistics

Results were reported as mean ± standard deviation (SD). Statistical analysis was performed using GraphPad Prism 9.0 (GraphPad Software, USA). Comparisons of experimental groups was performed with a one or two-way ANOVA, followed by post hoc Bonferroni correction. When normality could not be assumed, non-parametric tests were performed. Differences were found to be significant when * = p < 0.05, ** = p < 0.01, *** = p < 0.001, **** = p < 0.0001.

## Supporting Information

**Supporting Figure 1.**
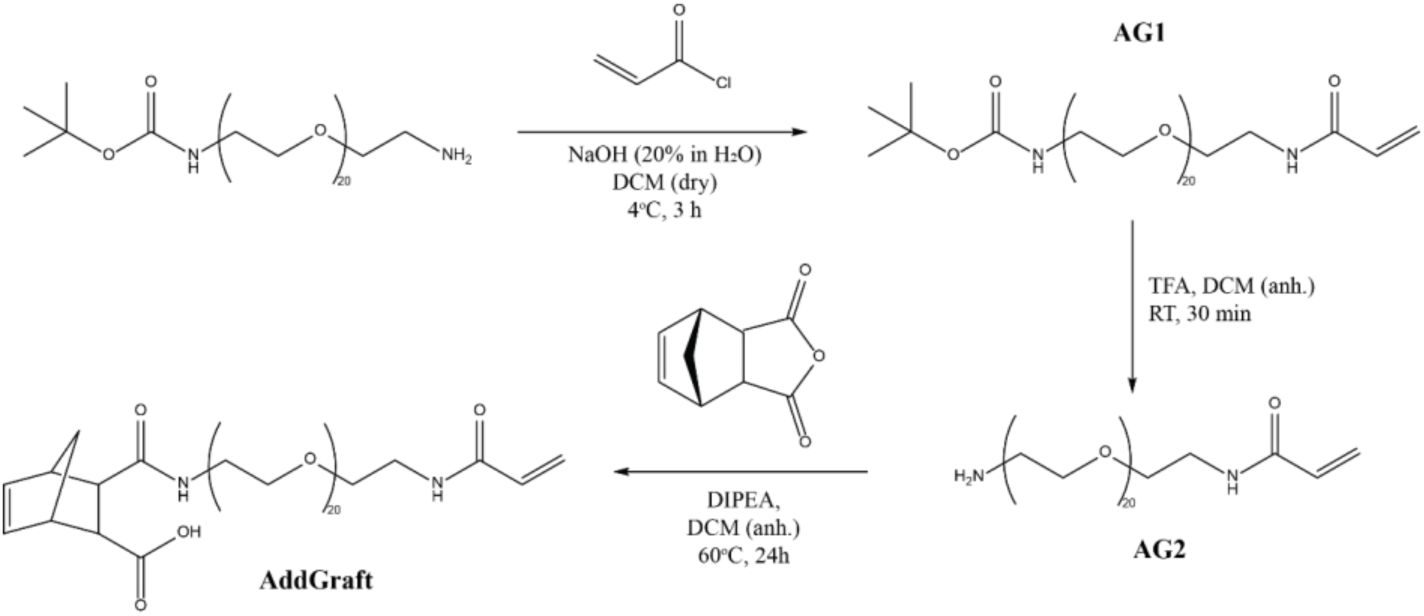
Schematic representation of the synthetic path to the complete synthesis of AddGraft.

**Supporting Figure 2.**
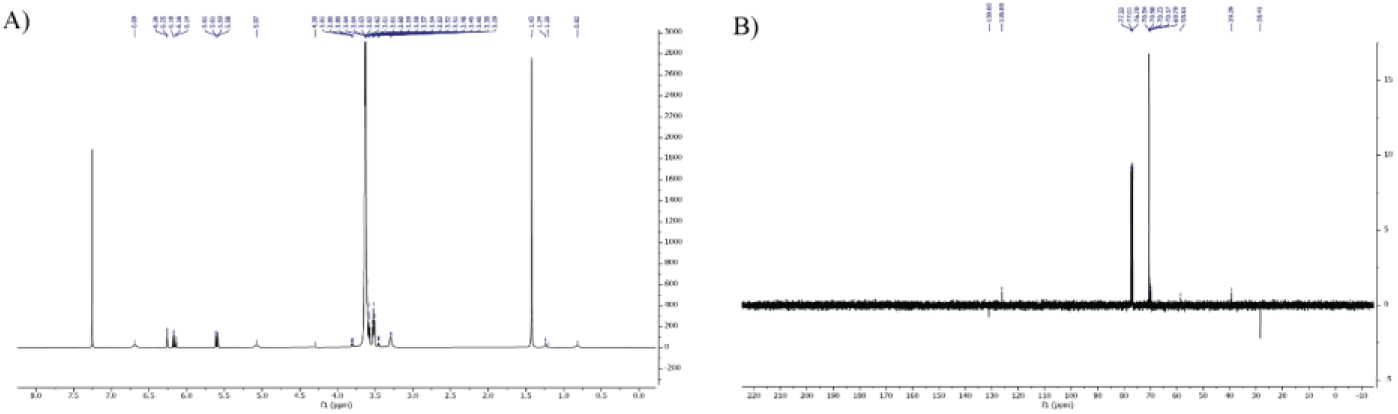
NMR characterization of AG1 in CDCl3. A) ^1^H NMR. B) ^13^C NMR.

**Supporting Figure 3.**
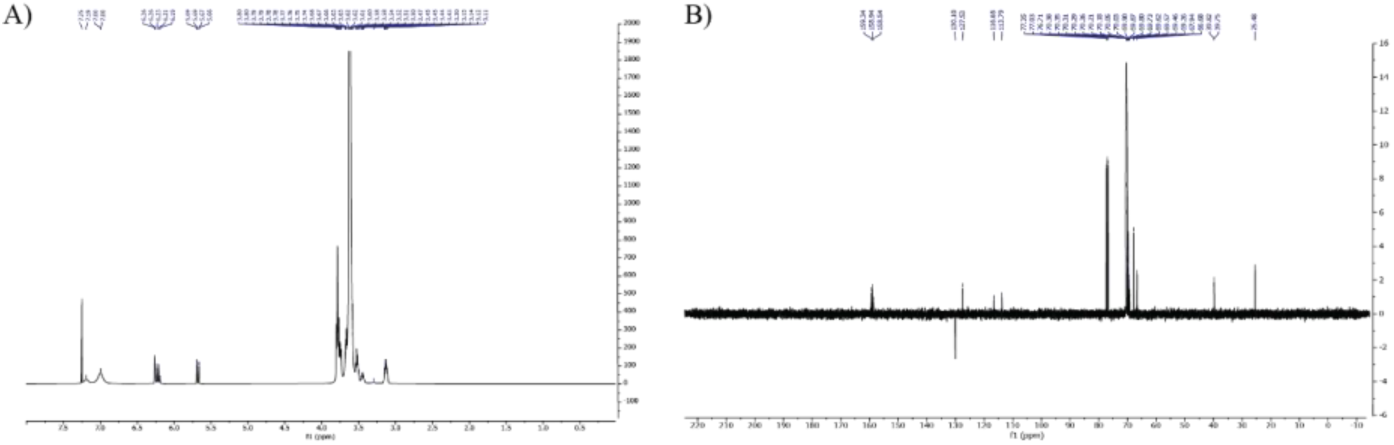
NMR characterization of AG2 in CDCl3. A) ^1^H NMR. B) ^13^C NMR.

**Supporting Figure 4.**
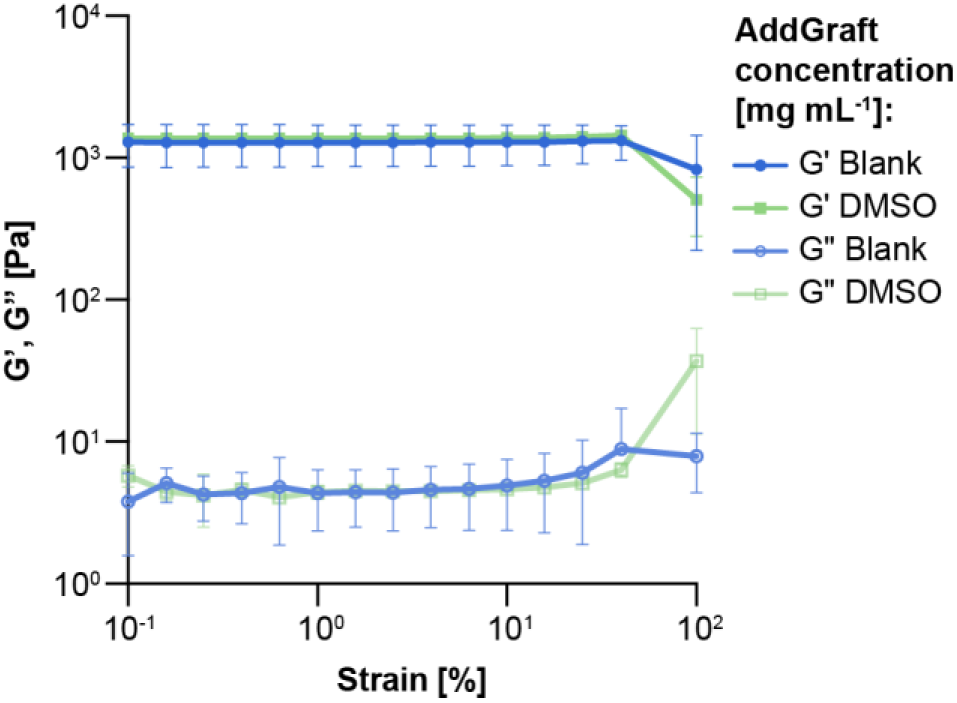
Photorheological amplitude sweep measurements of GelMA hydrogels with varying AddGraft concentrations displaying the linear viscoelastic range of the hydrogel material (n = 3).

**Supporting Figure 5.**
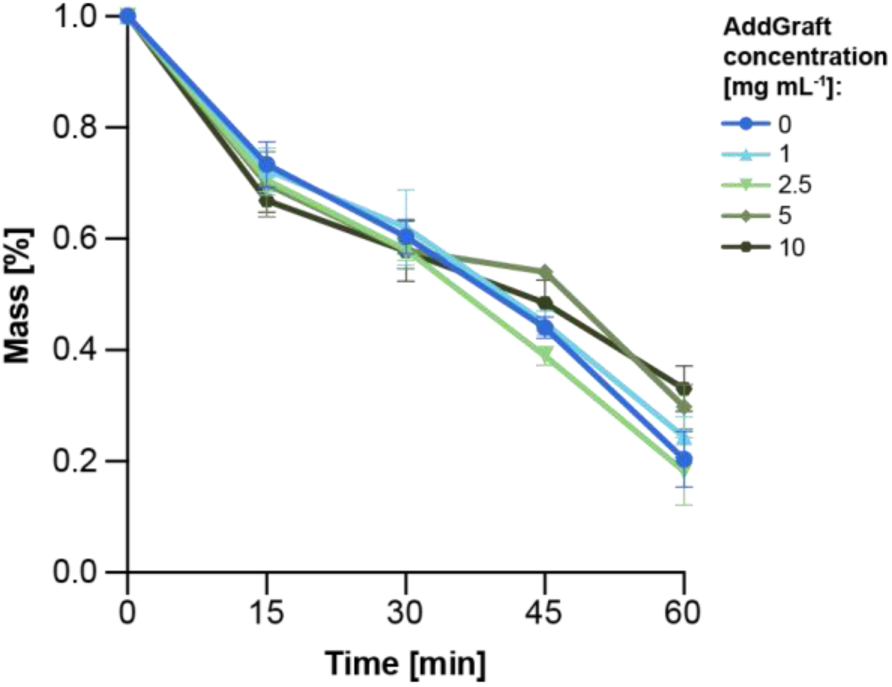
Enhanced enzymatic degradation assay of GelMA samples containing varying AddGraft concentrations by a 0.2 w/v% collagenase type II solution.

**Supporting Figure 6.**
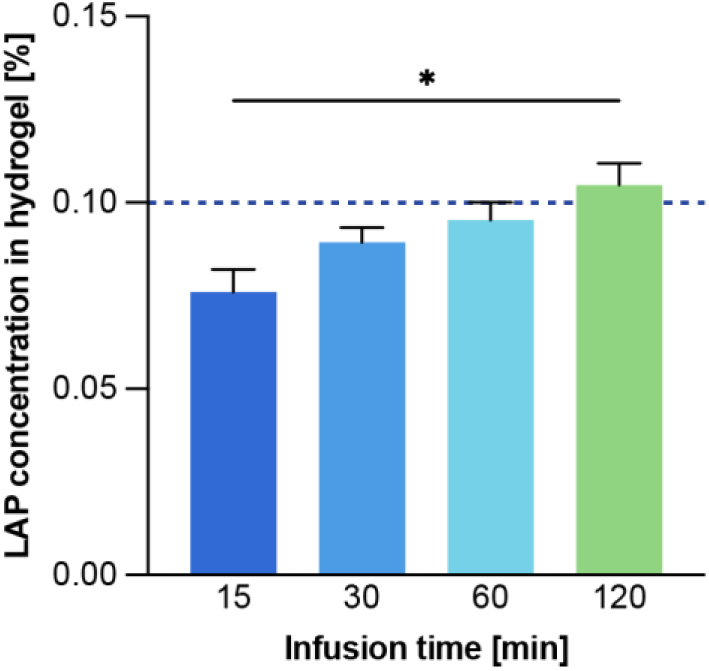
Analysis of LAP infusion rate into GelMA hydrogels.. A maximum of 0.1 w/v% LAP should physically be able to diffuse into the gel, displayed with a dotted line.

**Supporting Figure 7.**
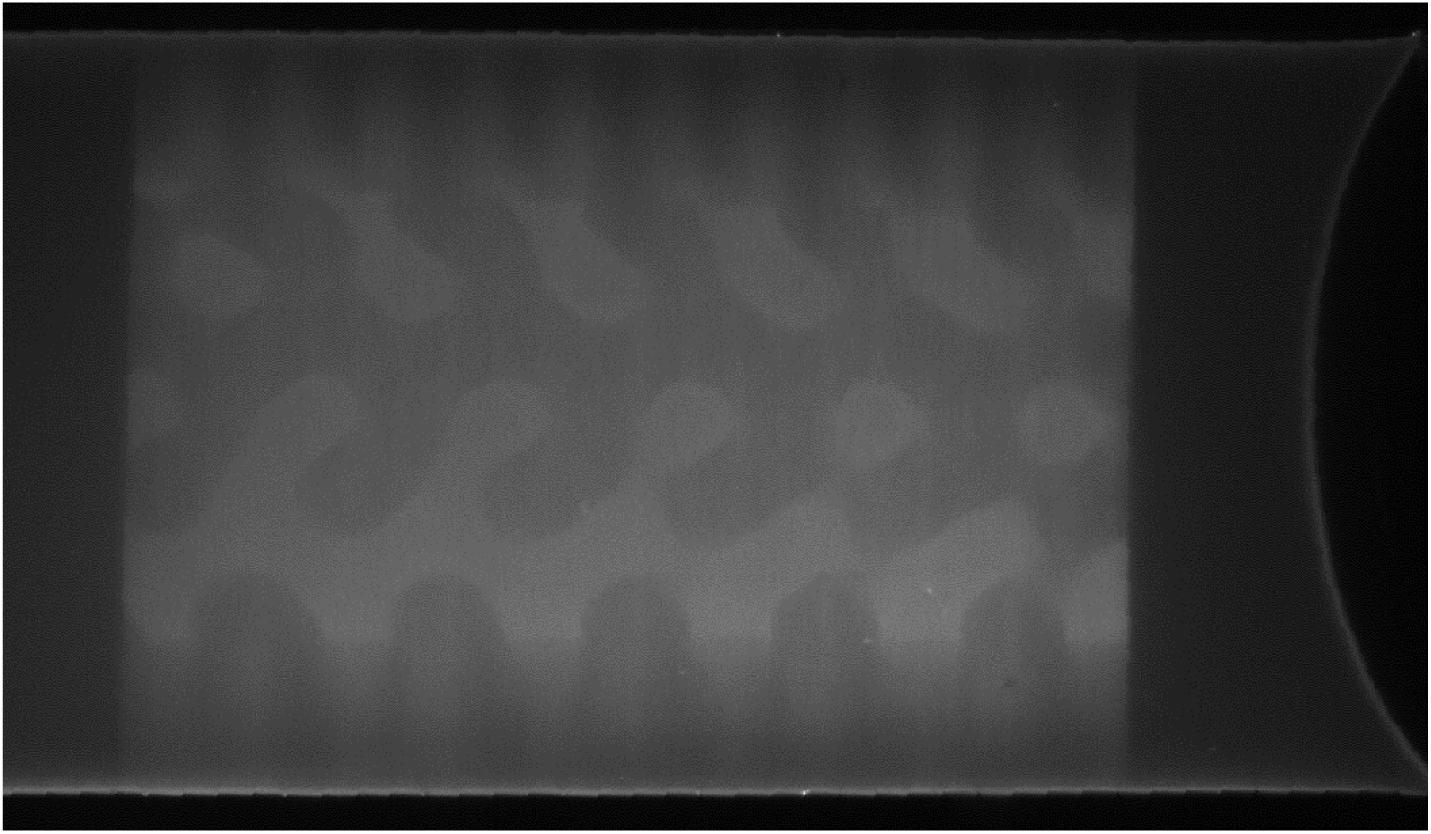
Z-slice image of lightsheet scan of a photopatterned gyroidal structure onto a AddGraft-containing GelMA hydrogel using Cy3-PEG5kDa-SH fluorescent dye.

**Supporting Figure 8.**
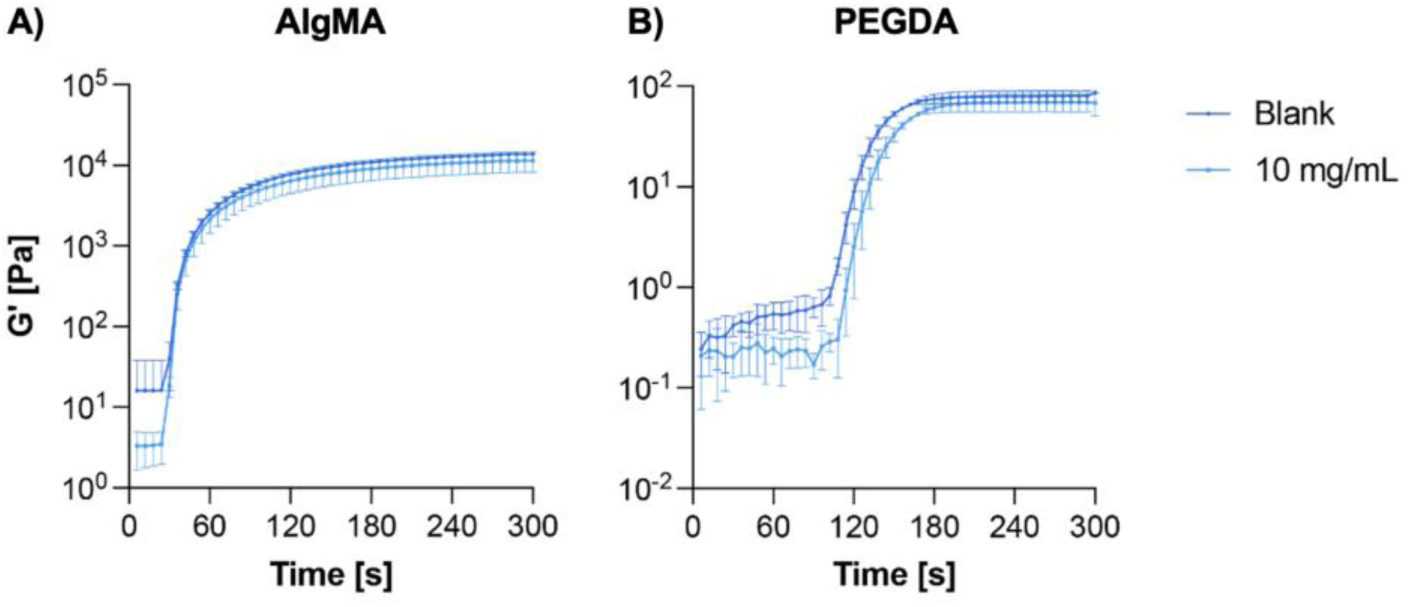
A) Photorheological time sweep of AlgMA hydrogels with and without AddGraft, displaying the photokinetics of the biomaterial with the addition of AddGraft (n = 3). B) Photorheological time sweep of PEGDA hydrogels with and without AddGraft, displaying the photokinetics of the biomaterial with the addition of AddGraft (n = 3).

**Supporting Figure 9.**
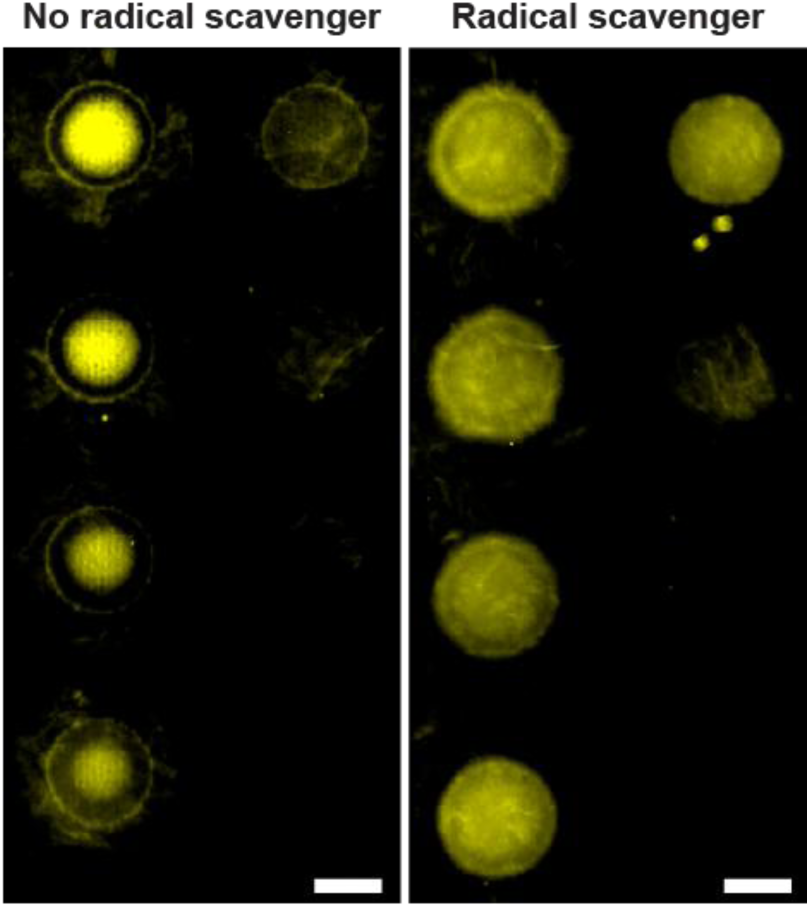
Representative dose test images of AlgMA hydrogels containing AddGraft with a varying TEMPO concentration and varying light dose range showing the effect of a radical inhibitor on the photobleaching of fluorescence.

**Supporting Figure 10.**
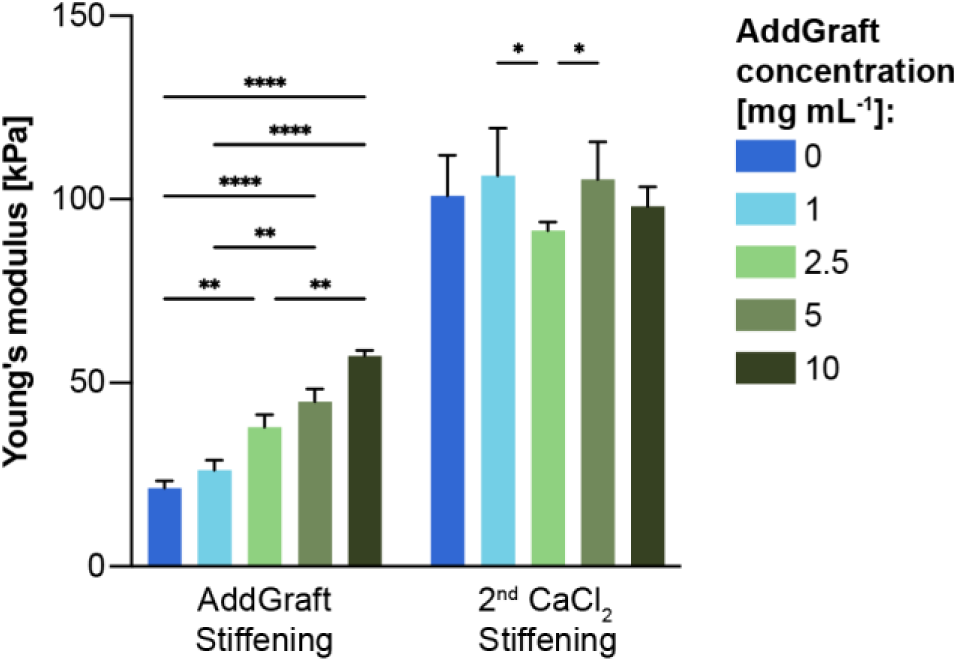
Compression modulus of AlgMA hydrogels with varying AddGraft concentrations after photostiffening and subsequently stiffened with a 50 mM CaCl2 solution.

**Supporting Figure 11.**
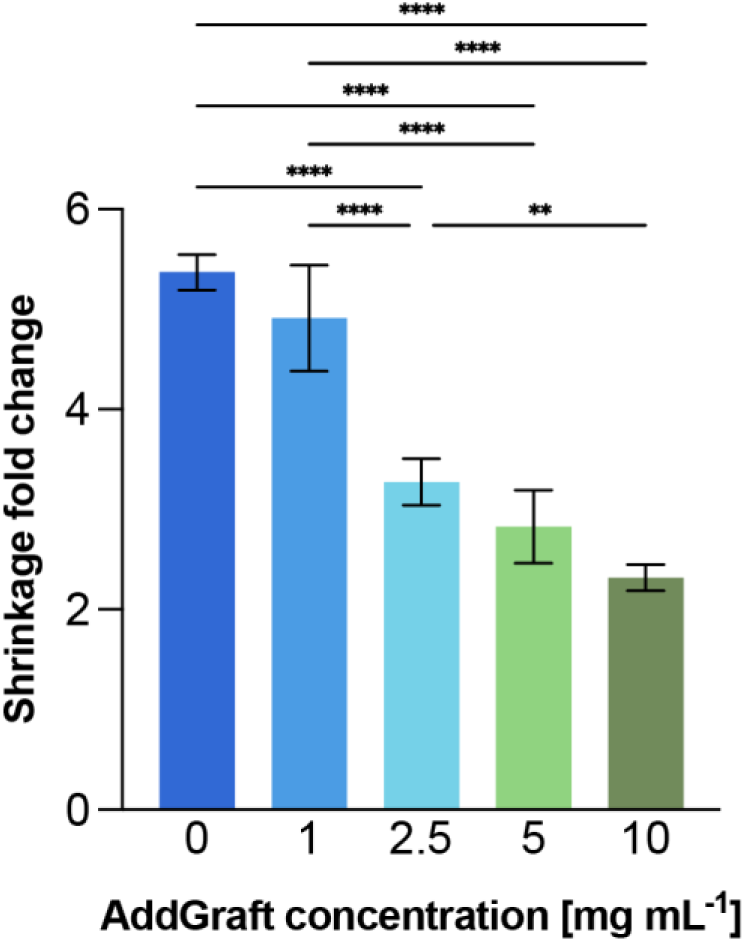
Shrinking analysis of AlgMA hydrogels with varying AddGraft concentrations after photostiffening and subsequently stiffened with a 50 mM CaCl2 solution.

**Supporting Figure 12.**
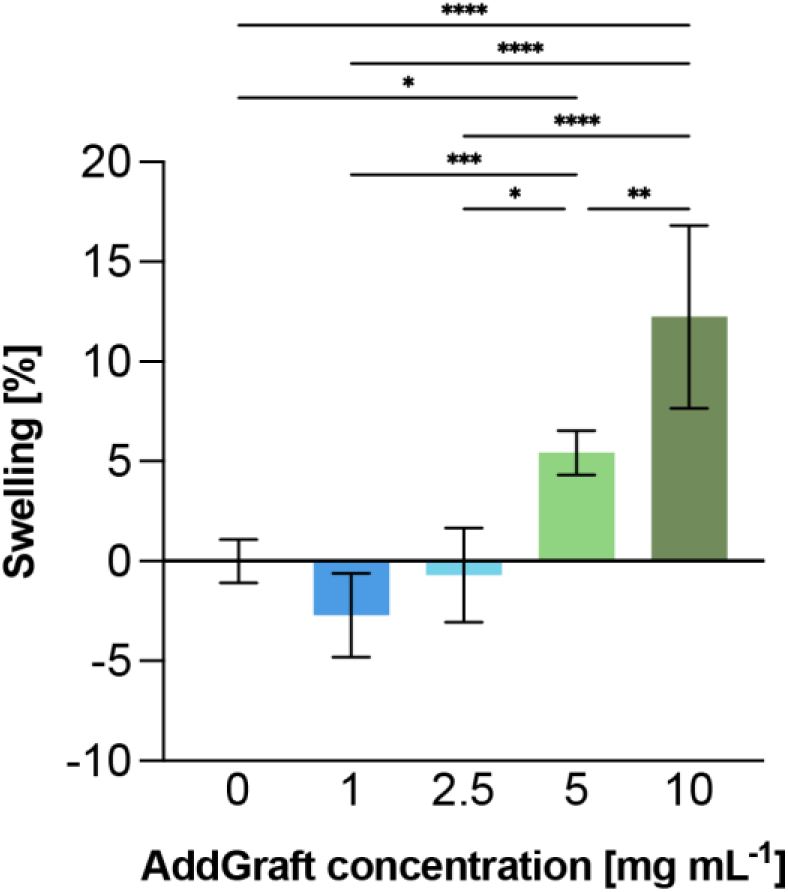
Swelling of PEGDA with varying AddGraft concentrations after photostiffening with 8-arm PEG thiol. Normalized to the control, no AddGraft, PEGDA.

**Supporting Figure 13.**
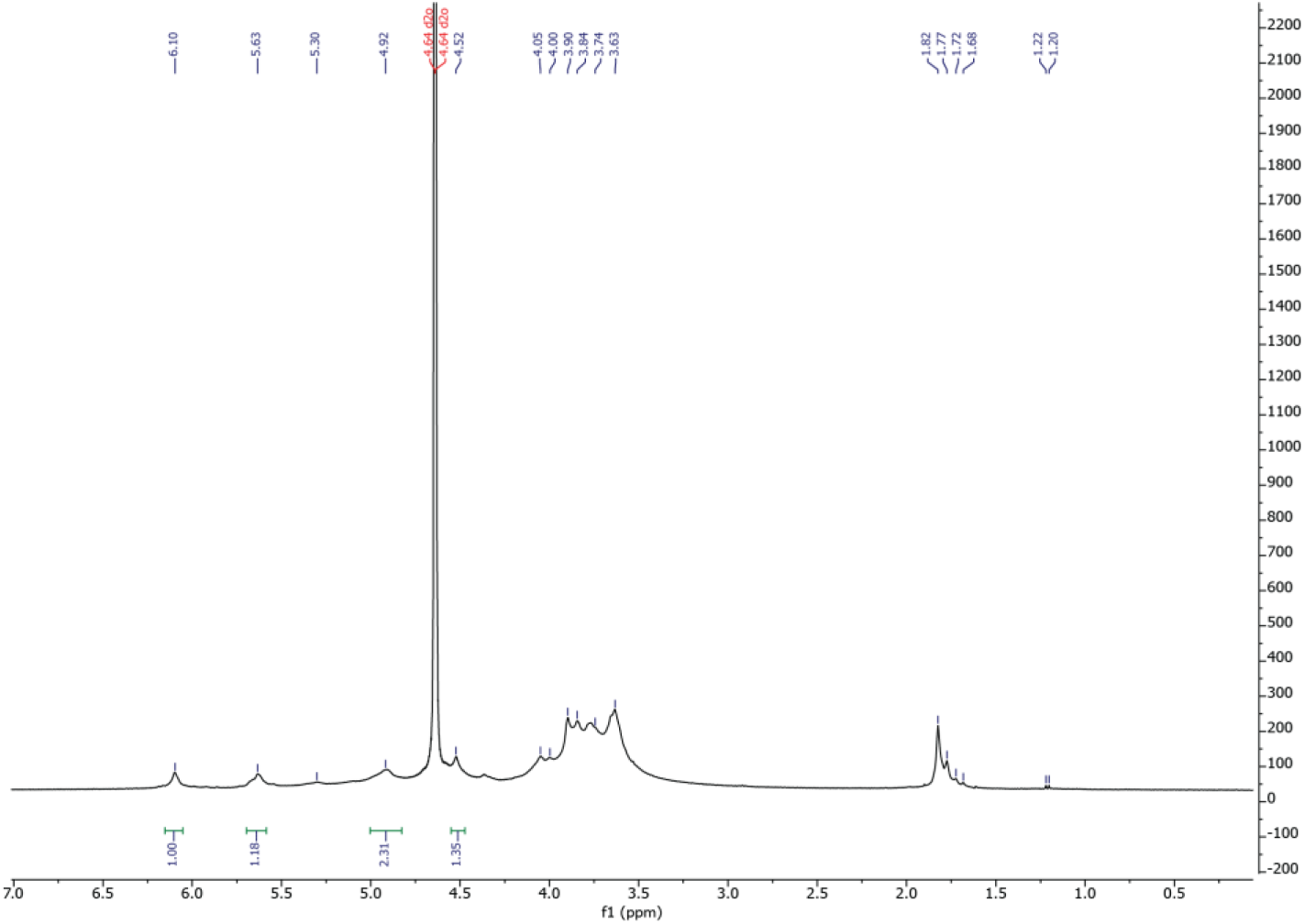
^1^H NMR of AlgMA for characterization of DoF.

